# Ketocarotenoid production in tomato triggers metabolic reprogramming and cellular adaptation: The quest for homeostasis?

**DOI:** 10.1101/2023.01.09.523254

**Authors:** Marilise Nogueira, Eugenia M. A. Enfissi, Elliott J. Price, Guillaume N. Menard, Eudri Venter, Peter J. Eastmond, Einat Bar, Efraim Lewinsohn, Paul D. Fraser

## Abstract

Plants are sessile and therefore have developed an extraordinary capacity to adapt to external signals. Here, the focus is on the plasticity of the plant cell to respond to new intracellular cues. Ketocarotenoids are high-value natural red pigments with potent antioxidant activity. In the present study, system level analyses have revealed that the heterologous biosynthesis of ketocarotenoids in tomato initiated a series of cellular and metabolic mechanisms to cope with the formation of metabolites that are non-endogenous to the plant. The broad multilevel changes were linked to, among others, (i) the remodelling of the plastidial membrane, where the synthesis and storage of ketocarotenoids occurs, (ii) the recruiting of core metabolic pathways for the generation of metabolite precursors and energy, and (iii) redox control. The role of the metabolites as regulators of cellular processes shown here, reinforces their pivotal role suggested in the remodelled “central dogma” concept.

## INTRODUCTION

The negative impact of climate change will continue to escalate with unchecked global CO_2_ emissions. Replacing petro-chemical derived products with plant biobased platforms will play an important role in achieving the Global Goals (UNEP, 2020). It is perceived that applied scientific endeavours will underpin this paradigm change to our manufacturing. However, the underlying progenitor to resolving these challenges are fundamental in nature. For example, to harness the full potential of plant-based systems, it is essential to define metabolic and cellular plasticity on a holistic level. Only through the acquisition of this fundamental knowledge will biosynthetic capacity extend to deliver biorenewables at commercially viable levels. Therefore, understanding how a plant cell adapts to metabolic perturbation and re-establishes homeostatic conditions is a fundamental aspect that underpins biotechnology in all its manifestations.

Ketocarotenoids are an example of high-value natural colorants and antioxidants used across multiple industrial sectors, including aquaculture where they are an essential feed additive. Although naturally present in some bacteria and algae, their present mode of industrial production uses chemical synthesis (Pfander et al., 1997). Total chemical synthesis used in this case has adverse environmental credentials, as the process uses petro-chemical derived precursors and rare metal catalysts. Hence, numerous examples exist whereby the formation of ketocarotenoids have been engineered into plant and microbial based systems (Rodriguez-Concepcion et al., 2018). The generation of a high ketocarotenoid tomato line and demonstration of its production, technical and economic feasibility is an exemplar of a sustainable plant-based renewable source of ketocarotenoids for aquaculture (Nogueira et al., 2017). It was estimated that all the ketocarotenoid feed supplements required annually for aquaculture could be generated from this ketocarotenoid tomato genotype grown in 15,000 ha of greenhouse (Napier et al., 2020).

Beyond its value as a biobased platform, the ketocarotenoid tomato is a fundamental resource for the study of cell plasticity, whereby complex biochemical and cellular adaptations in response to non-endogenous metabolite production can be revealed. In the present study, the ketocarotenoid line has been characterised along with a wild type and orange fruited segregants using an integrated multi-omic approach. This systems approach revealed the diverse range of adaptation mechanisms harnessed by the plant cell to facilitate the heterologous biosynthesis and storage of ketocarotenoids in the plastid membranes without inducing pleiotropic effects. Homeostasis is described as the self-regulatory processes deployed by an organism, tissue or cell to maintain internal stability in response to external changes (Billman, 2020). Here, we discuss this concept in the context of internal changes occurring from the production of non-endogenous molecules in cells.

Furthermore, these findings have been put into perspective with respect to the fundamental knowledge of cellular flux. The original central dogma of molecular biology, first presented by Francis Crick in 1957, described the transfer of information during DNA replication, transcription into RNA and translation into protein (Crick, 1970, Crick, 1958). For decades, metabolism has been overlooked in the central dogma (Schreiber, 2005) until advances in metabolomics shed light on the role of metabolites as fine and coarse regulators of biological systems (Kochanowski et al., 2017, Yang et al., 2018, Luca et al., 2020). This prompted the central dogma concept to be reviewed (Costa Dos Santos et al., 2021). This paper supports the pivotal role of metabolites in the remodelled central dogma concept.

Altogether, this study provides fundamental knowledge across the different levels of cell regulation and has direct impact on the exploitation of future biotechnological platforms for the creation of a bioeconomy.

## RESULTS

The ketocarotenoid line was previously generated by crossing a low producing ketocarotenoid line (ZW), overexpressing the *Brevundimonas* β-carotene hydroxylase (*CrtZ*) and β-carotene ketolase (*CrtW*) (Enfissi et al., 2019) with an orange fruited recombinant inbred line accumulating a high level of β-carotene, the main precursor of the ketocarotenoid biosynthetic pathway (Nogueira et al., 2017). In this study, the ketocarotenoid line was compared to the β-carotene line (orange fruited segregant), which resulted from the segregation of the *CrtZ* and *CrtW* genes and to the control line which is the wild type segregant.

### Changes in chlorophylls, carotenoids and their esterification in the engineered tomato lines

The ketocarotenoid line, the β-carotene line and the control line, had very distinct fruit phenotypes which reflected their carotenoid composition (Figure 1a). From 49 days post anthesis (dpa), the red fruit of the control line and the orange fruit of the β-carotene line predominantly accumulated lycopene (red pigment) and β-carotene (orange pigment), respectively (Figure 1a & Table S1). In tomato fruit, the lycopene cyclase enzyme (CYC-B) is responsible for the conversion of lycopene to β-carotene (Figure 1b). As described previously (Alba et al., 2005, Ronen et al., 2000), its expression decreased at 43 dpa, which resulted in an accumulation of lycopene in the control fruit (Figure 1a). However, in the β-carotene line, the presence of a highly expressed fruit ripening enhanced lycopene cyclase gene (*Cyc-b*-Gal, ∼15 fold increase compared to the control *Cyc-b* at 49 dpa, Figure S1), led to high levels of β-carotene (Figure 1a). From early development (25 dpa), the ketocarotenoid line accumulated a complex mixture of ketocarotenoids (deep red pigments), in free and esterified forms (Figure 1a & S2). Total ketocarotenoid content reached 2.5 mg/g DW in the 66 dpa keto fruit (Figure 1a, Table S1). The constant presence of a range of ketocarotenoids is explained by the constitutive expression of the heterologous *CrtZW* genes and the partial conversion of ketocarotenoid precursors to astaxanthin, the end product of this pathway (Figure 1b & S2). Only ketocarotenoids harbouring a hydroxyl moiety, such as adonixanthin, phoenicoxanthin and astaxanthin can be esterified (Figure 1b). Those ketocarotenoids have all been found in their esterified form in the fruit (Figure 1a & S2, Table S1). The predominant ketocarotenoid, phoenicoxanthin, accounted for almost half of the ketocarotenoids quantified at 66 dpa. Esterification of phoenicoxanthin increased over fruit development from approximatively 10% to 50% (Figure 1a & S2, Table S1). The ketocarotenoid fruit also differed by its chlorophyll content, which decreased (2 to 3-fold) compared to the control and the β-carotene line (Figure S3). The constitutive expression of *CrtZW* resulted in the presence of ketocarotenoids in the flowers and leaves of the ketocarotenoid line, leading to observable orange coloured petal and dark green to brown leaf hue (Figure 1a). Similarly to the fruit, in the leaf, a complex ketocarotenoid profile was observed and the chlorophyll content was decreased (∼1.5-fold) compared to that in the control and β-carotene lines (Figure S4a). Moreover, the efficiency of the photosystem II, which is reflected by the Fv/Fm value (the maximum photochemical efficiency of PSII in the dark-adapted state), was shown to be negatively affected (16% significant reduction, Figure S4b). Furthermore, as expected in leaf since the *Cyc-b*-Gal expression is fruit ripening enhanced, the production of ketocarotenoids depleted endogenous precursors, such as β-carotene (2.5-fold lower relative to controls) as well as other endogenous carotenoids (Figure S4, Figure 1).

**Figure 1:**
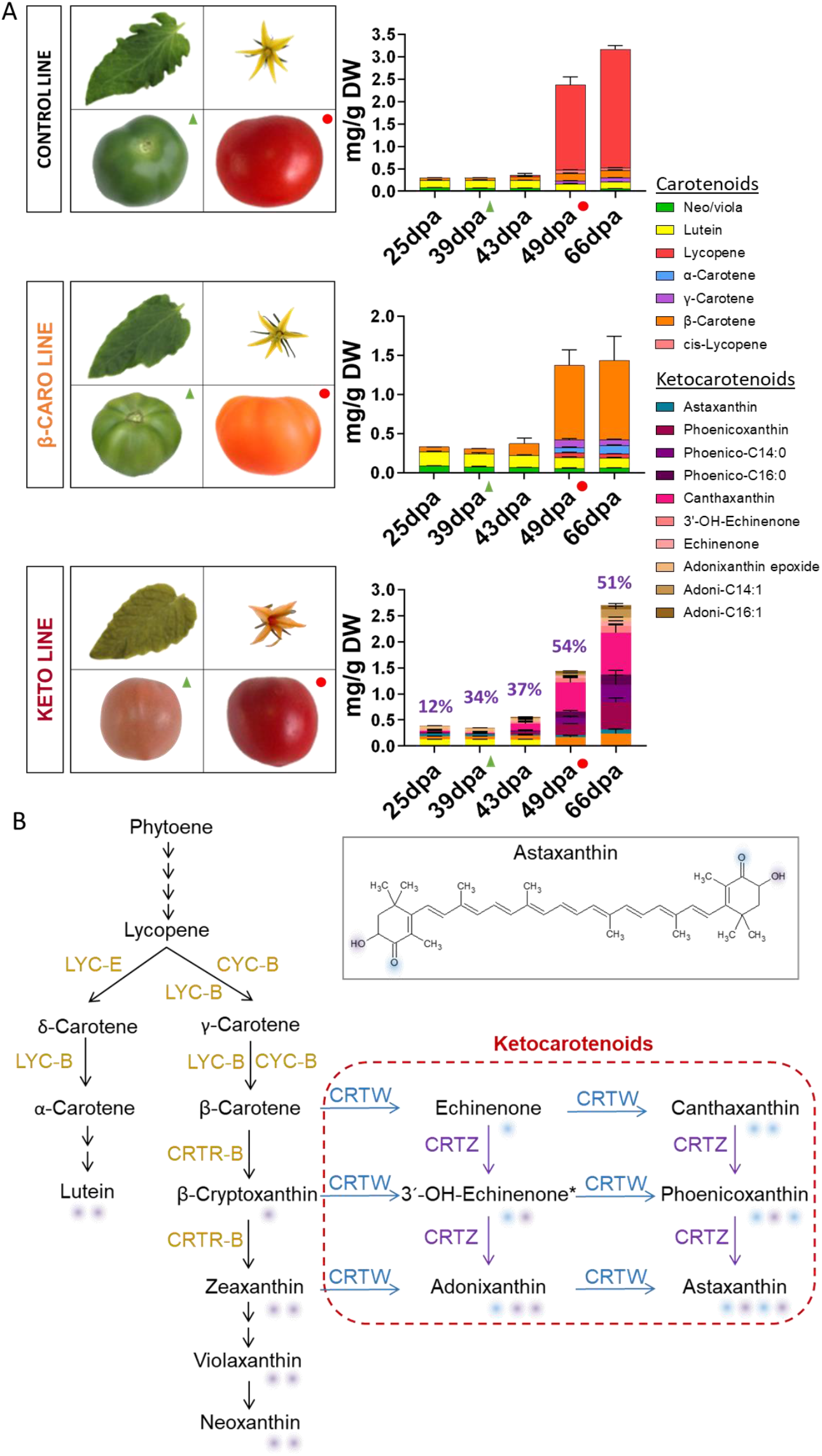
Tomato plant phenotypes and keto/carotenoid composition at five fruit ripening stages. (A) Phenotype of the leaf, flower, mature green fruit (39 dpa) and ripe fruit (49 dpa) of the control, β-carotene (β-caro) and ketocarotenoid (keto) lines and their fruit keto/carotenoid composition (only pigmented compounds are displayed) at five ripening stages (25, 39 (green triangle), 43, 49 (red circle), 66 days post anthesis). Data represent mean in mg/g dry weight ± SD. The rate of esterification of the main ketocarotenoid, phoenicoxanthin, is shown as % in the graph of the keto line. Astaxanthin esters data were not included; Neo, neoxanthin; Viola, violaxanthin; Phoenico, phoenicoxanthin; Adoni, adonixanthin. (B) Biosynthetic pathway of the endogenous carotenoids and heterologous ketocarotenoids (red rectangle) introduced in tomato plant. Enzyme names are as follow: LCY-E, ε-lycopene cyclase; LCY-B, β-lycopene cyclase; CYC-B, fruit ripening enhanced β-lycopene cyclase; CRTR-B1, carotene β-hydroxylase 1; CRTW, bacterial carotene ketolase and CRTZ, bacterial carotene hydroxylase (Brevundimonas sp.). The purple and blue shadings depict the position of the functional hydroxyl and ketone groups, respectively

### Ultrastructural changes to plastid structures resulting from altered carotenoid content

Electron micrographs (EM) of the plastids were analysed to study the structural impact of the synthesis and storage of the ketocarotenoids on the cell. At mature green stage, the chloroplasts of the different lines looked similar, all presenting characteristic chloroplast sub-compartments (thylakoid membranes and small plastoglobules). However, some differences were observed in the chloroplasts from the keto line compared to those of the other lines. The number of vesicles, which are derived from the chloroplast inner envelope membrane (Nogueira et al., 2013), were increased in the keto chloroplasts (Figure 2a). Almost 50% of the keto EM showed chloroplasts containing more than 10 vesicles compared to only 6% of the control EM (Table S2a). At ripe stage, as expected, the chloroplasts were converted to chromoplasts, the thylakoid membranes disintegrated and the plastoglobules were of greater size. The chromoplasts of the different lines showed striking changes (Figure 2a). The plastoglobules in the keto chromoplasts represented a significant greater area (1.6 μm^2^) compared to the ones in the control line (0.3 μm^2^). The difference was due to an increased plastid area as well as a greater plastoglobule number in the keto chromoplasts (Table S2b). The plastoglobules of the keto chromoplasts also displayed a significant darker colour intensity (grey ratio of 4.4) compared to the plastoglobules of the other lines (grey ratio of 2.2). The same trend was observed at mature green (Table S2b). Compounds, such as plastoquinone and α-tocopherol, are expected to contribute to the osmiophilicity of the plastoglobules (electron-opaque/darkness) thanks to their ethylenic double bonds which reduce the TEM staining agent OsO4 (van Wijk and Kessler, 2017, Merriam, 1958). Both compounds were found in significantly greater levels in the keto chromoplasts compared to their levels in the control and β-carotene chromoplasts (Figure 2b). Consequently, their increased contents correlated with the darker plastoglobules observed in the EM.

**Figure 2:**
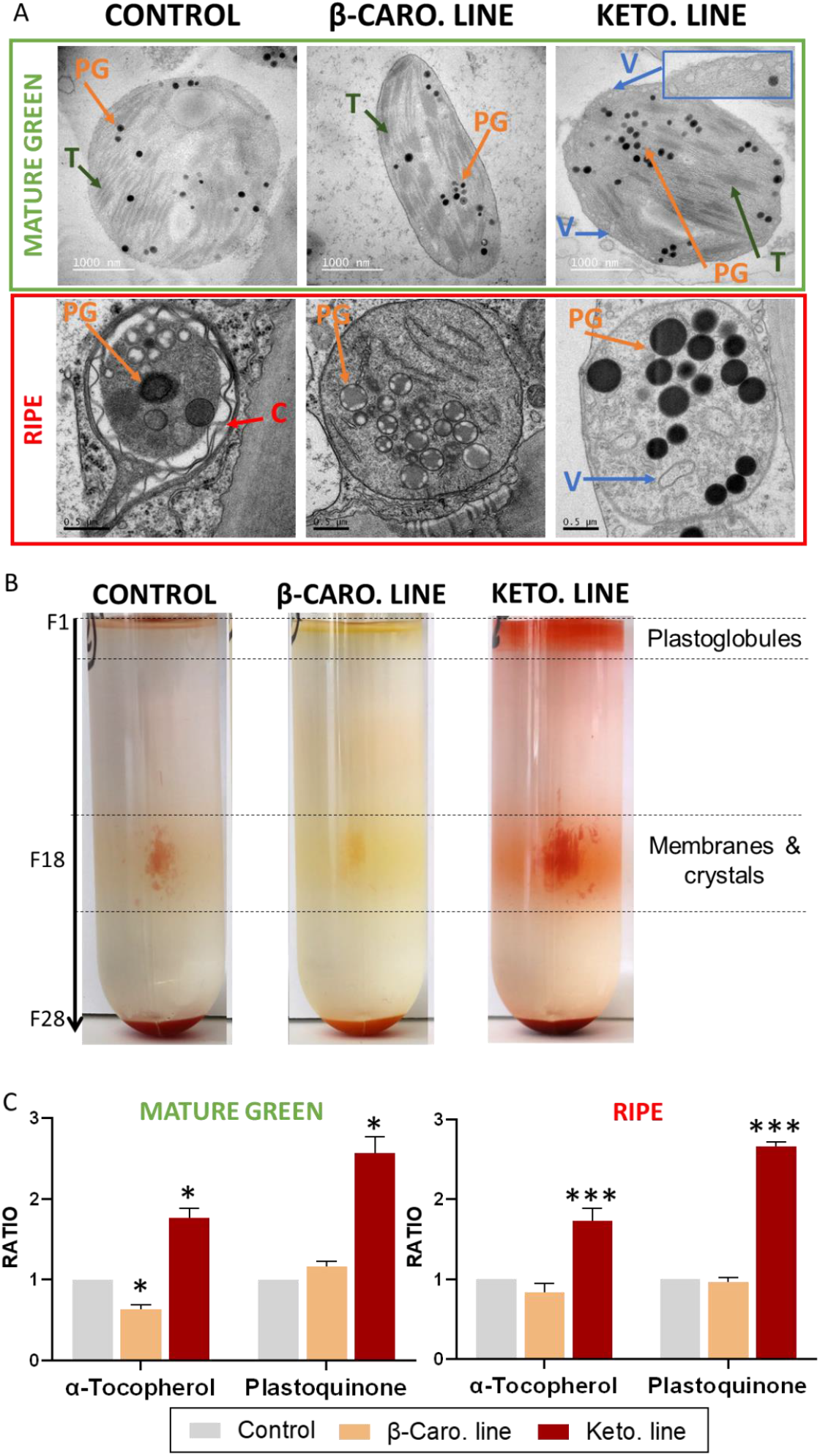
Ketocarotenoid sequestration and its effect on subplastidal compartment structures. (A) Electron micrographs of chloroplasts (Mature green fruits, 39 dpa) and chromoplasts (Ripe fruits, 49 dpa) of the control, β-carotene (β-caro) and ketocarotenoid (keto) lines. T, thylakoid; PG, plastoglobule; V, vesicles also called membranous sacs; C, lycopene crystal. The blue insert is a zoom in (2X) on the chloroplast envelope and its vesicles. (B) Fractionation of subchromoplast compartments of red fruit (∼ 47 dpa) of the control, β-carotene and ketocarotenoid lines. Each fraction (F) corresponds to 1 ml. (C) Alpha-tocopherol and plastoquinone contents in the mature green and ripe fruit of the control, β-carotene and ketocarotenoid lines. Data represent the ratio of the means compared to the control mean ± SD. Asterisks indicate significant differences compared to the control line (ANOVA analysis). P < 0.05, P < 0.01, and P < 0.001 are designated by *, **, and ***, respectively.

To investigate the subplastidial localisation of the ketocarotenoids in the different lines, sub-chromoplast compartments were separated on a sucrose gradient. Ketocarotenoids were found in the membranes, mainly in crystalline form, and in the plastoglobules of the keto chromoplasts, while the predominant carotenoids in the other lines, lycopene and β-carotene were observed in the membranes and some also in crystalline form (Figure 2c, Table S3). It is worth noting that although β-carotene is a cyclic carotenoid like the ketocarotenoids, only ketocarotenoids were stored in the plastoglobules (Figure S5). The esterified ketocarotenoids, like their free form, were also found in both the membranes and plastoglobule fractions (Figure S5). The (keto)carotenoids (i.e. carotenoids and ketocarotenoids), which contain unsaturated double bonds, are also expected to affect plastoglobule osmiophilicity. Therefore, the ketocarotenoid localisation also correlated with the darker plastoglobules observed in the EM of the keto chromoplasts.

### Holistic changes to metabolism beyond the formation of carotenoids

To study how the cell metabolism adapted in response to the ketocarotenoid production, a wide range of metabolites (amino acids, sugars, organic acids, carboxylic acids, phytohormones, phenolics, phytosterols, fatty acids, volatiles, etc.) were analysed (Table S4, S5). A Principal Component Analysis (PCA) integrating all metabolite data measured in ripe tomatoes, except the (keto)carotenoids data, revealed that, all the biological replicates of each line formed independent clusters and that the greatest metabolic difference was observed between the keto line and the other lines (Figure S6). This result was representative of the PCA plots obtained for each analytical platform separately (i.e. LC-QQQ-MS, LC-QTOF-MS, GC-MS, SPME-GC-MS, UPLC-DAD). This highlighted that the alteration of the keto line metabolism affected compounds beyond the carotenoid pathway and that the variation between the keto line metabolism and the other lines was greater than the variation observed between the metabolism of the control and β-carotene lines.

The analysis of catabolism revealed striking differences in the volatile content of the ripe fruit from different lines (Figure 3a, Table S4). Each line contained carotenoid derived volatiles which corresponded to the cleavage of predominant (keto)carotenoid(s) in the fruit. For instance, (i) β-cyclocitral and β-ionone, β-carotene derived volatiles, were found in proportionally greater amounts in the β-carotene line (ii) sulcatone and epoxygeraniol, lycopene derived volatiles, in the control line and (iii) oxo-β-cyclocitral and oxo-β-ionone (non-endogenous) in the keto line. Some volatile levels were increased up to 20-fold and oxo-β-ionone, was uniquely observed in the keto line (Figure 3b). A PCA of the volatile data revealed that the three tomato lines were not only characterised by their respective (keto)carotenoid volatiles but also, (i) for the β-carotene line, by its phenolic/shikimate and monoterpene derived compounds and (ii) for the ketocarotenoid line, by the branched-chain amino acid (leucine and valine) derived volatiles. The composition of fatty acid derived compounds was different in all three lines. It is interesting to note that, penten-3-one, which derives from the C18:3 fatty acid, was increased by 2.5-fold in the keto line (Table S4).

**Figure 3:**
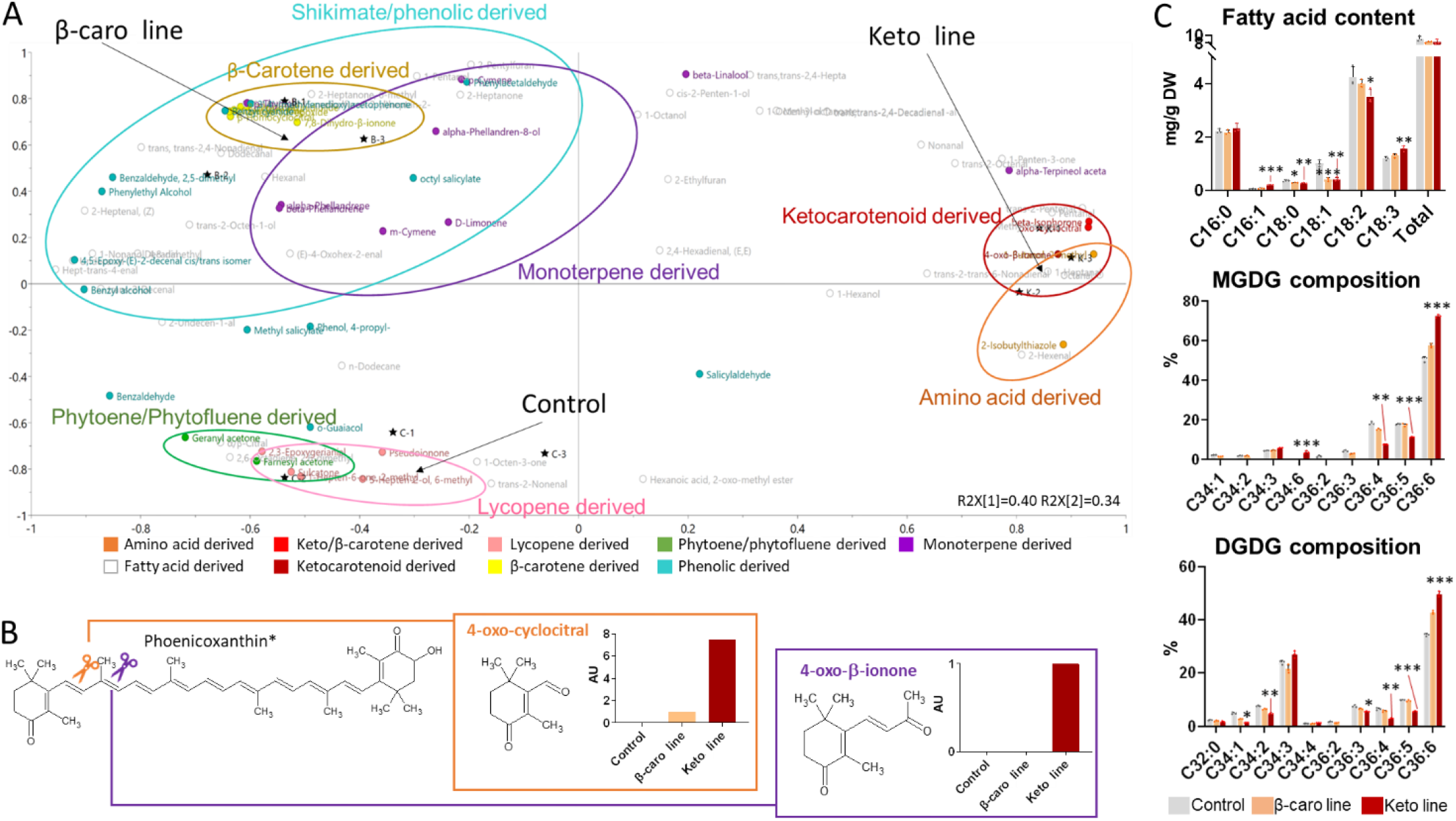
Effect of ketocarotenoid production on volatile and fatty acid contents in ripe tomato fruit. (A) Loading plots of principal component analysis of volatiles measured in ripe fruit of the control, β-carotene (β-caro) line and/or ketocarotenoid (keto) line. Volatile data are listed in Table S4. (B) Formation of the ketocarotenoid derived volatiles. The abundance of the volatiles in the control, β-carotene line and/or ketocarotenoid line is displayed in bar charts; AU, arbitrary unit; *, phoenicoxanthin or any 4-oxo ketocarotenoid. (C) Fatty acid content and plastidial lipid (MGDG, DGDG) fatty acid composition in fruit. Data represent mean in mg/g dry weight ± SD. Asterisks indicate significant differences compared to the control line (ANOVA analysis). P < 0.05, P < 0.01, and P < 0.001 are designated by *, **, and ***, respectively. P-values are listed in Table S13. MGDG, Monogalactosyldiacylglycerol; DGDG, digalactosyldiacylglycerol.

The alteration of the volatile profiles suggested changes in the cellular metabolism therefore the fatty acid content in the ripe tomato was examined. The total fatty acid content was not significantly different between the three lines however the fatty acid composition of the ripe keto tomato was shifted towards the synthesis of fatty acid with increased unsaturation (Figure 3c). Indeed, the content of C18:0, C18:1 and C18:2 was significantly decreased while C18:3 level was increased by 1.3-fold compared to that in the control line (Table S4). Moreover, an increase of the C16:1 content (3-fold) was also observed in the keto line. Similarly, the fatty acid composition of the MGDG and DGDG lipids, which are structural lipids of the plastid membranes, also showed an increased in the percentage of C36:6 (C18:3+C18:3) at the expense of the less unsaturated lipids C36:3 to C36:5 (Figure 3c). It is interesting to note that C34:6 (C18:3+C16:3) was only detected in the MGDG analysis of the keto tomatoes.

Ratios of metabolite levels in the keto line relative to the control were displayed on a pathway (Figure 4). The keto ripe fruit contained decreased levels of sugars such as sucrose, fructose and galactose and increased levels of amino acids (valine, cysteine and serine), phytosterols (campesterol, stigmasterol and β-sitosterol) and phenolics (naringenin, chlorogenic acid and feruloyquininc acid). The levels of some unknown phenolics, such as the unknown designated unk_m/z:432.2, were increased by more than 100-fold in the keto line. The changes in volatile contents often correlated with those of the metabolite levels from which they were derived. For instance, the levels of valine and valine derived volatiles were both increased in the keto line. Minor changes in phytohormone levels were observed such as, at MG, an increase in abscisic acid and at ripe, an increase in jasmonic acid and a decrease in auxin. The levels of TCA cycle metabolites were consistently altered at MG and ripe, with an accumulation of citric acid, aconitic acid and mesaconic acid and a decrease of succinic acid, fumaric acid and malic acid (Figure 4, Table S4, S5).

**Figure 4:**
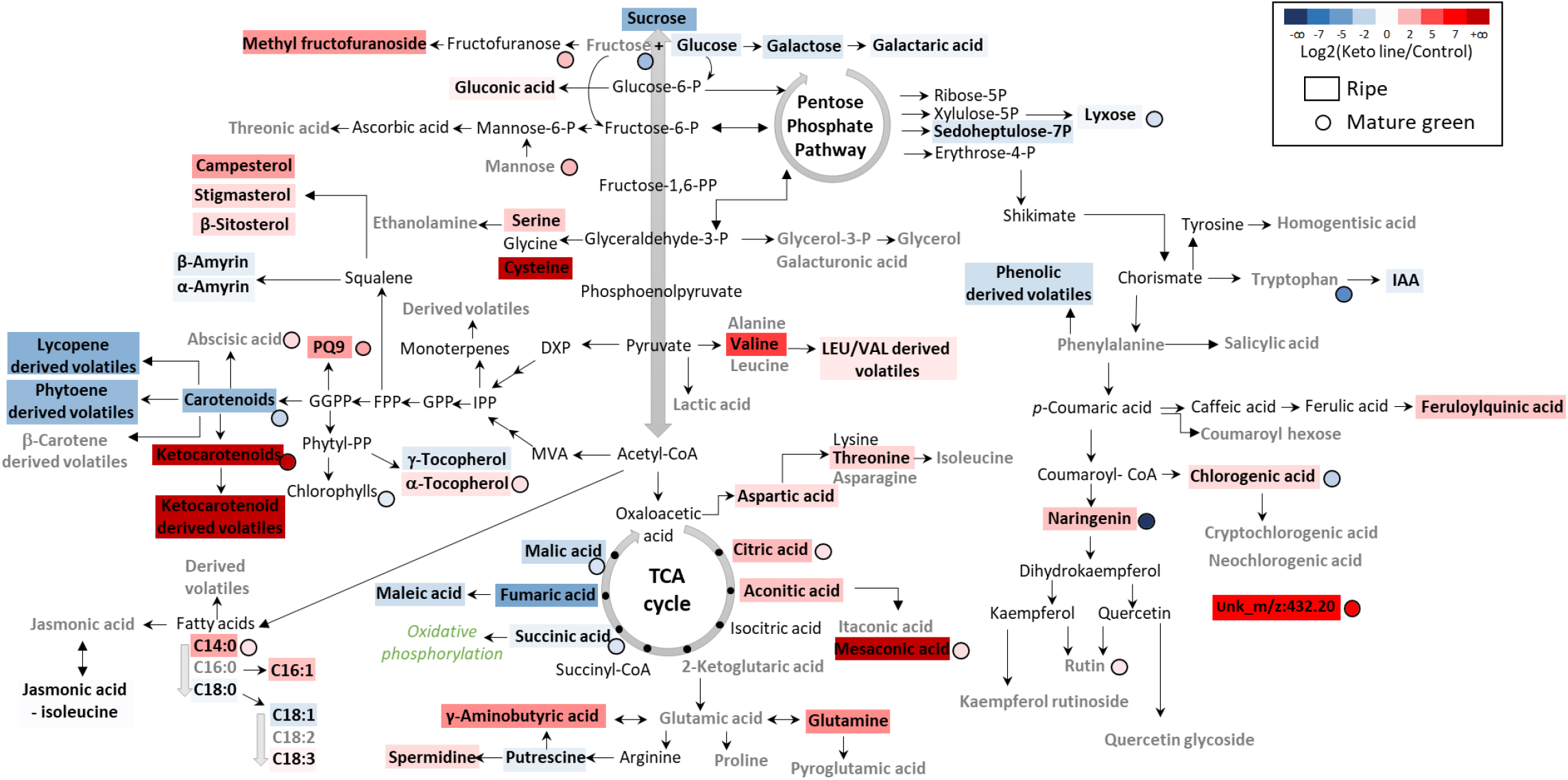
Biochemical pathways displaying the fold change metabolite levels in ripe and mature green fruit of the ketocarotenoid line compared to the control. Data represent the log2 of the ratio of means (Keto line/Control) and it is displayed by a colour illustrating a decrease (blue) or increase (red) of metabolite level in the keto line compared to the control. Compounds detected in the analytical platforms are shown in bold. The absence of significant difference is depicted by a grey bold font (t-test, P>0.05). Compounds showing a significant difference (black bold font) were attributed a colour representative of fold change. Metabolite levels quantified in ripe and mature green fruits are represented by a rectangle and a circle, respectively. The statistical significance of the volatile data in this display corresponds to the total content per class of compounds. The individual differences within the class of compounds are not shown. Metabolite data were compiled in Table S4 and S5. P-values are listed in Table S13. PQ9, plastoquinone 9; IAA, Indole-3-acetic acid

The high production of β-carotene in tomato fruit also led to global metabolic changes (Figure S7). Similarly to differences observed in the keto ripe fruit, the levels of some TCA cycle metabolites were also different at both MG and ripe in the β-caro fruit compared to the control (decrease of malic acid and fumaric acid levels). However, in general, fewer pathways were altered compared to those in the keto/control line comparison (Figure 4, Figure S7).

Comparably to the ripe stage, the control, β-caro and keto lines could also be differentiated by their metabolic profiles at MG independently of their carotenoid content (Figure S8a). The metabolic profile of the keto fruit was characterised by its content in carboxylic acids and phytohormones, while the profile of the β-caro fruit was distinguishable by its elevated amino acid content and high levels of jasmonic acid and jasmonic acid-isoleucine, and finally the control line was differentiated by its phenolic and fatty acid contents (Figure S8b, Table S5).

### Impact of the ketocarotenoid production at a transcriptomic level

The variation of the transcriptome between the ketocarotenoid line, β-carotene line and control line was investigated in MG and ripe fruit. Thereafter, the differentially expressed genes (DEG) obtained from the RNA seq comparison of the keto and the β-carotene lines compared to the control line are referred to as keto/control and β-carotene/control DEG, respectively. In general, the numbers of DEG found at the ripe stage were greater compared to the ones at MG for all DEG sets (Figure 5a, Figure S9a). The higher number of DEG in the ripe fruit was concurrent with the significant accumulation of the (keto)carotenoids (Figure 1a). Moreover, at both developmental stages, the keto/control DEG were greater, in terms of number, compared to the β-carotene/control ones, 1834 vs 1152 at ripe and 1200 vs 988 at MG, respectively. This is reflected in the RNAseq dendrogram of the ripe fruit, which showed that the keto line was the most distinct compared to the other lines (Figure S10). Therefore, both transcriptomic and metabolomic data (Figure S10 & S6b) indicated that the alterations in the ripe keto fruit were the greatest compared to the control. At MG, β-carotene/control DEG were evident and corroborated the findings from metabolomic analysis (Figure 5a, Figure S7).

**Figure 5:**
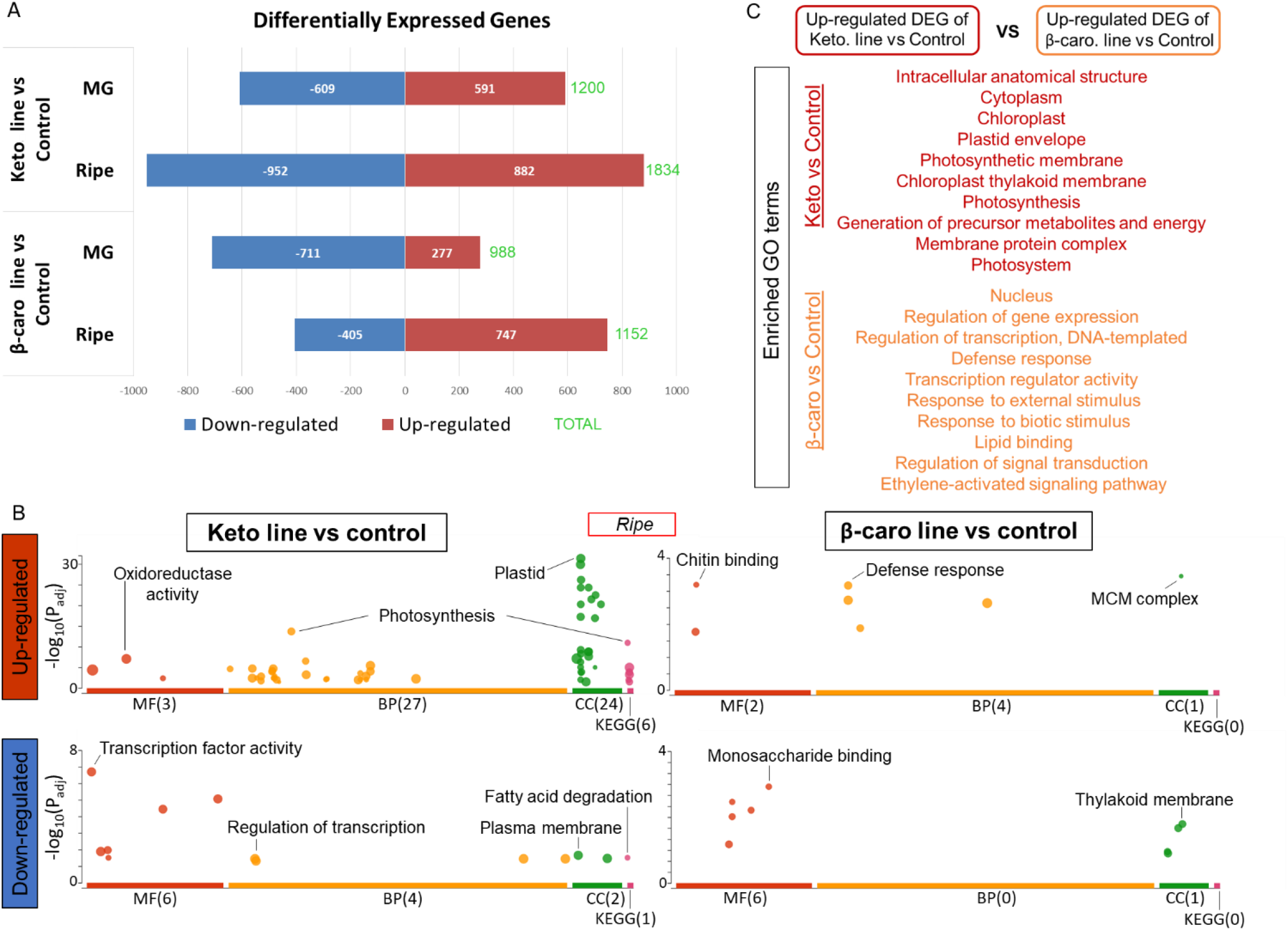
Differentially expressed genes and enrichment analysis of the transcriptomic data comparison of the tomato lines. (A) Up-regulated and down-regulated differentially expressed genes (DEG) between the ketocarotenoid and β-carotene lines compared to the control, respectively, at mature green (MG) and ripe fruit stage. (B) Enriched Gene Ontology (GO) terms of the up-regulated and down-regulated DEG sets of the ketocarotenoid and β-carotene lines versus control comparison, respectively. MF, molecular function; BP, biological process; CC, cellular component; KEGG, KEGG pathway. (C) Gene enrichment analysis of the up-regulated DEG obtained from the ketocarotenoid line versus control comparison with the up-regulated DEG from the β-carotene line versus control comparison. The full list of enriched GO terms can be found in Table S11.

Gene ontology (GO) enrichment analyses confirmed that, when ripe, the keto/control comparison harboured up-regulated DEG with many more enriched GO terms compared to those in the β-carotene/control DEG (54 vs 7, respectively) and also compared to those found in the down-regulated DEG (12 and 7, respectively, Figure 5b). In the ripe fruit, the enriched GO terms of the up-regulated keto/control DEG were related to plastid membranes, photosynthesis, the generation of precursor metabolites and energy, oxidoreductase activity, carbon metabolism, biosynthesis of secondary metabolites and response to hydrogen peroxide. The enriched GO terms in the down-regulated keto/control DEG were mainly linked to transcription regulation and fatty acid degradation (Figure 5b, Table S6). At MG, a greater number of enriched GO terms were found in the down-regulated keto/control and β-carotene/control DEG (35 and 54, respectively) compared to those in the up-regulated DEG analyses (9 and 4, respectively). The enriched GO terms of the down-regulated keto/control DEG at MG were related to the photosynthetic electron transport chain and the cell wall, while the ones found in the up-regulated keto/control DEG were linked, for instance, to the response to ABA and to oxygen containing compounds (Figure S11, Table S6).

The transcriptomic data of the ripe fruit have been displayed over biosynthetic pathways in Figure 6 (Details in Table S7). The keto/control comparison revealed that genes of the carotenoid pathway were up-regulated in the keto line, as well as genes encoding biosynthetic steps in associated pathways such as GGPP formation, supplying both the tocopherol and plastoquinone pathways (Figure 6). Up-stream pathways providing carotenoid precursors, such as the MEP pathway, were also up-regulated. Furthermore, core metabolic pathways such as the glycolysis, the TCA cycle, the pentose phosphate pathway and the photosynthesis, photorespiration, chromorespiration as well as electron transport chain were also positively regulated. Perturbations to the fatty acid and lipid biosynthetic pathway as well as phenolic formation were also observed. Therefore, the transcription has been altered not only for the carotenoid biosynthetic genes or closely related metabolic pathway genes but also at a global level whereby core metabolic pathway gene transcription was altered (Figure 6). The transcriptomic data of the β-carotene/control comparison highlighted the up-regulation of the transcription in the carotenoid pathway and down-stream pathways such as ABA formation (Figure S12 & Table S7). Alteration of transcription was also observed within the phenolic pathway and among core metabolic pathways but to a lesser extent compared to those seen in the keto/control comparison. Moreover, in the β-carotene line, most changes in the core pathways were mainly down-regulation of gene expression.

**Figure 6:**
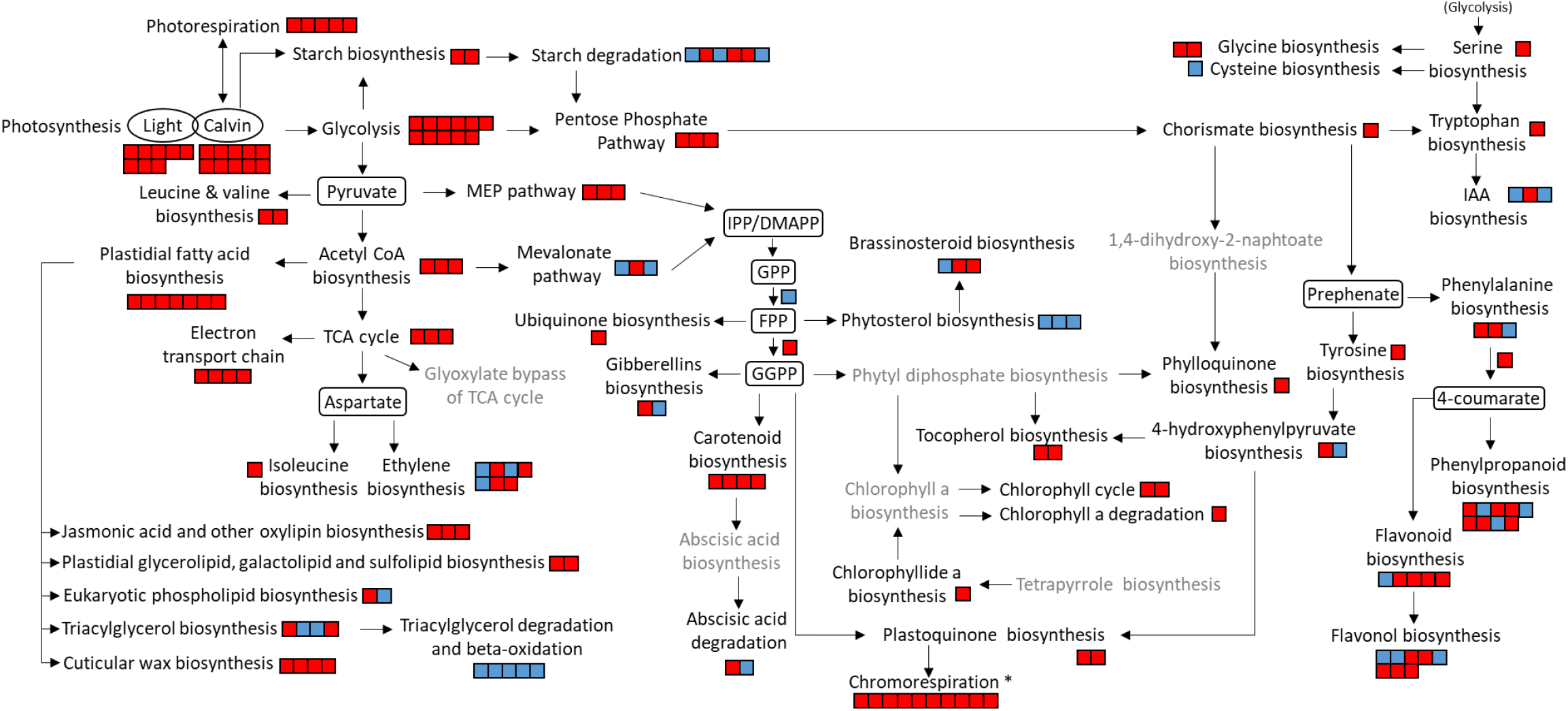
Transcriptional changes in the metabolic pathways of the ripe fruit of the keto line compared to the control. RNAseq data was used to create this display. Each square represents a gene of the referred pathway. Red and blue colours illustrate an increase and decrease in expression, respectively (positive or negative log2(Fold change)). The grey font denotes a lack of significant difference in expression level for all the genes investigated in the respective pathway (P > 0.05). All genes used to construct this figure are listed in Table S7 together with their log2(Fold change). Gramene database (www.gramene.org) as well as literature were used to compose the different metabolic pathways. IPP, Isopentenyl pyrophosphate; DMAPP, Dimethylallyl pyrophosphate; GPP, Geranyl pyrophosphate; FPP, Farnesyl diphosphate; GGPP, Geranylgeranyl pyrophosphate ; IAA, Indole-3-acetic acid ; TCA, Tricarboxylic acid cycle ; * according to models proposed by Renato et al., 2014 and Grabsztunowicz et al., 2019.

The endogenous β-carotene level in the keto fruit seemed to have been maintained thanks to the changes in *Cyc-B*-Gal gene expression. Indeed, *Cyc-B*-Gal was up-regulated by ∼15-fold in the ripe keto and the β-carotene fruits compared to the control fruit and by ∼4-fold at MG but only in the keto fruit (Figure S1). On the other hand, the expression of the chloroplast lycopene cyclase *Lcy-b*, which is responsible for the formation of β-carotene in the MG fruit, was unchanged compared to the control in both tomato lines, and at both developmental stages studied (Figure S1). It was demonstrated earlier that the esterification of ketocarotenoids occurred from the early fruit developmental stage. The transcriptomic data revealed that, *gdsl esterase* expression was up-regulated (2.6-fold) at MG while *pyp-1 esterase* expression was up-regulated (2-fold) at ripe compared to the control. PYP1 enzyme has been shown to esterify ketocarotenoids in tomato fruit (Lewis et al., 2021). Altogether, it shows that the adaptation to the presence of ketocarotenoids was fluid and evolving through time.

The chloroplastic malate dehydrogenase enzyme is considered a marker for the plastidial redox status (Nashilevitz et al., 2010, Obata, 2015). Its expression was up-regulated (1.6-fold) in the keto line compared to that in the control (Table S8). Similarly, many other genes related to redox control, such as thioredoxin, ferredoxin, cys-peroxiredoxin, the chloroplastic cytochrome b6, were also up-regulated (up to 3-fold). In particular, the expression of the plastidial NAD(P)H-quinone oxidoreductase subunit M (Ndh-M) was increased by 20 and 11-fold in the ripe keto fruit compared to that in the control and β-carotene lines, respectively (Table S8). Ndh-M protein is part of the NAD(P)H dehydrogenase complex (Ndh), which supports non-photochemical electron fluxes from stromal electron donors to plastoquinones. In the chromoplast, this respiratory process can be referred to as chromorespiration, which is thought to take place within the vesicles or membranous sacs of the chromoplasts (Renato et al., 2014). The Ndh complex activity was proven to link redox activity to central and specialised metabolism such as carotenoid, tocopherol and phenolic metabolism (Nashilevitz et al., 2010). Here, a network analysis of the transcriptome and metabolome data of the keto/control comparison revealed that *Ndh-M* was positively correlated to ketocarotenoids (p-corr = 0.97 and 0.94) and to the content of total phenolics (p-corr = 0.7) (Figure 7, Table S9). Moreover, one of the ketocarotenoids, phoenicoxanthin-C16:0, was found to be positively correlated to, among others, *Ndh-M* (p-corr = 0.97), *rubisco* (p-corr = 0.96), *phenylalanine ammonia-lyase* (p-corr = 0.96), and *α-amylase* (p-corr = 0.97) genes. This reinforces the idea that the ketocarotenoid biosynthesis is linked with core and specialised metabolism but also to redox control. Additionally, the expression of other Ndh subunits (B, L, N) were also all up-regulated (up to 7-fold) in the keto line (Table S8). Similarly, the expression of the gene coding for PsbQ-like protein, which is required for the chloroplastic Ndh complex function (Yabuta et al., 2010) increased 15-fold in the keto line compared to the control (Table S8).

**Figure 7:**
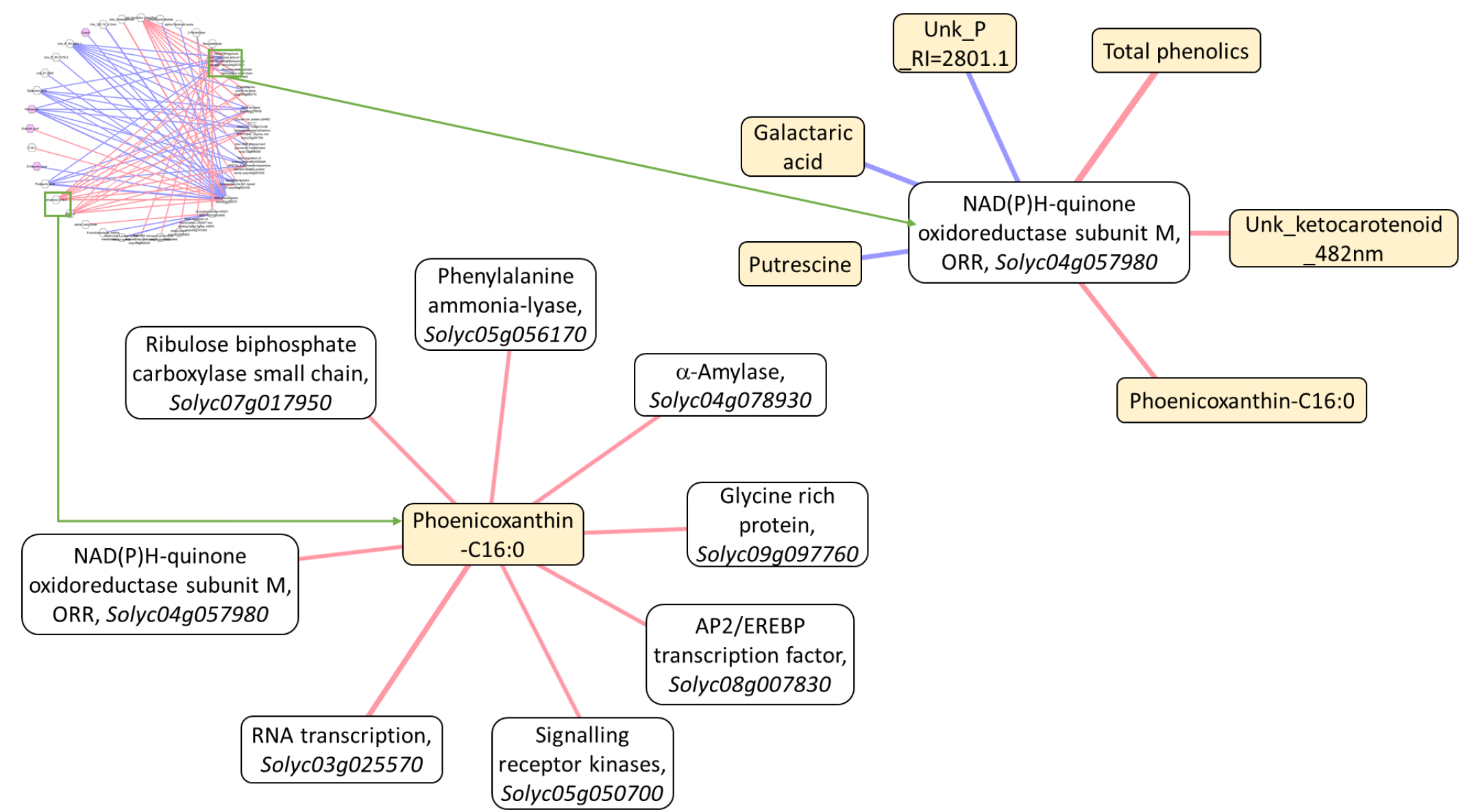
Gene-metabolite correlation network analysis of the ripe keto line /control comparison. Metabolites and genes related to biosynthetic pathways, whose levels showed a statistical difference in the keto line compared to the control (p< 0.05), were selected to create the correlation network. Sub-networks involving the ketocarotenoid phoenicoxanthin-C16:0, and the gene NAD(P)H-quinone oxidoreductase subunit M are shown separately. Metabolites and genes are represented in yellow and white rounded rectangle, respectively. Positive and negative correlations are denoted as pink and purple edges, respectively. The relative width of edges corresponds to the absolute value of the correlation coefficient. The Person correlation values are given in Table S9.

In order to study the differences between the keto line and the control over ripening, GO enrichment analysis was carried out comparing the respective Ripe/MG DEG sets. It revealed that significantly less genes related to the GO terms “plastid” and “photosynthesis” had their expression level decreased over ripening in the keto line compared to those in the control, suggesting that plastid metabolism is, at least partially, kept on in the keto line over ripening (Table S10).

Finally, to investigate how the cell, in the ripe fruit, was adapting differently when faced with the biosynthesis of the non-endogenous ketocarotenoids compared to the biosynthesis of the endogenous β-carotene, GO enrichment analysis was performed comparing the keto/control and β-carotene/control up-regulated DEG sets at ripe stage (Figure 5c, Table S11). The enriched GO terms for the adaptation to the ketocarotenoid production were linked to the plastid membranes, the photosynthesis and the generation of precursor metabolites and energy, while the enriched GO terms for the adaption to β-carotene production were mainly related to regulation of gene expression, regulation of primary metabolic process and response to stimulus. Similarly, to understand the characteristic responses caused by the formation and storage of the ketocarotenoids, GO enrichment analysis was carried out on the common keto/control and the keto/β-carotene DEGs at ripe (Figure 8). The enriched GO terms in the common up-regulated DEGs were associated with the plastid, photosynthesis, generation of precursor metabolite and energy and oxidoreductase activity, while those representing the down-regulated DEG were connected to the nucleus, the transcription and regulation mechanisms of biosynthetic processes (Table S12).

**Figure 8:**
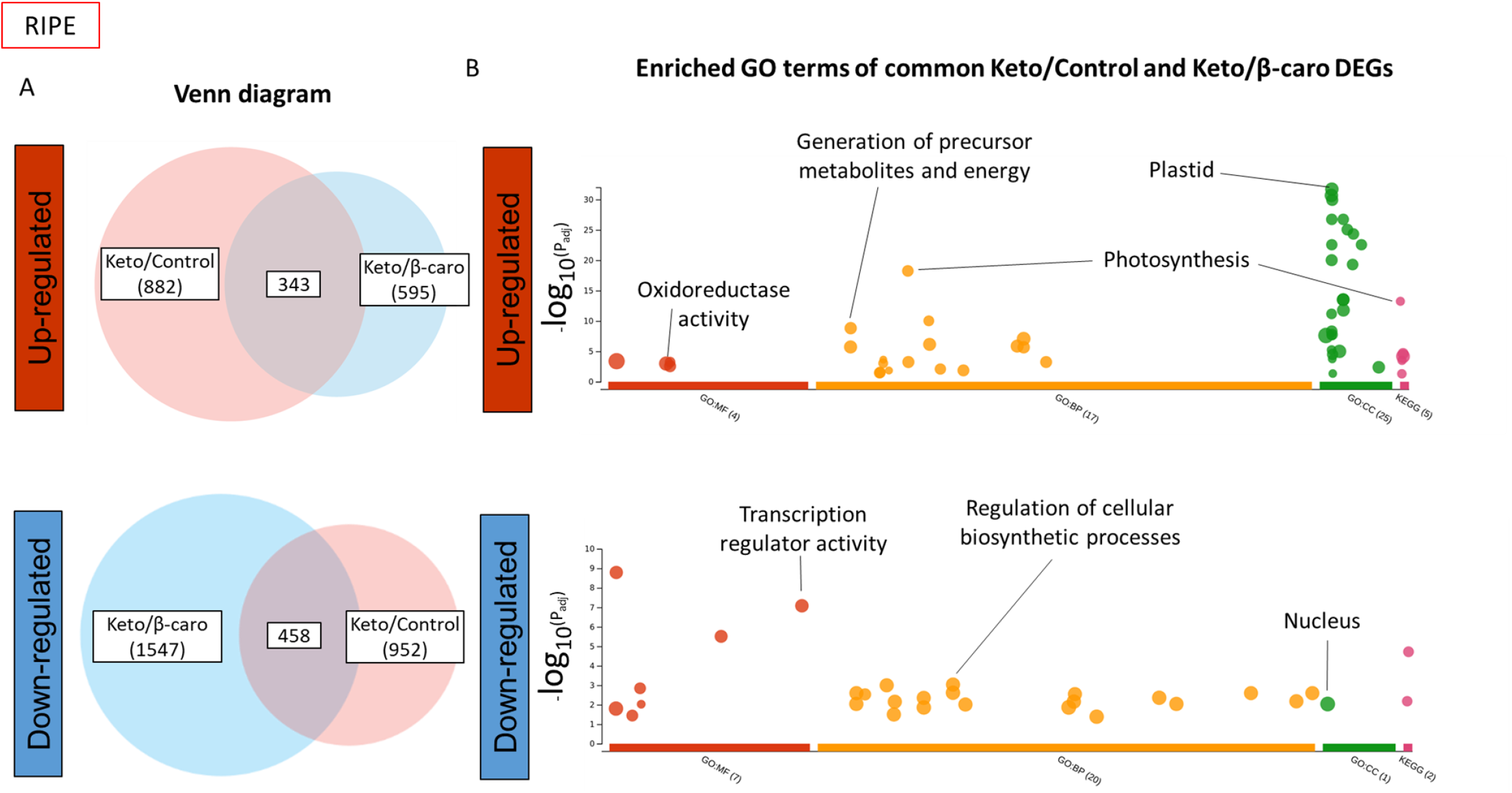
Gene enrichment analysis of the common DEGS of Keto/Control and Keto/β-caro sets in ripe fruit. (A) Venn diagrams of the up-regulated and down-regulated DEGs of the Keto/Control and Keto/β-caro sets (B) Enriched GO terms of the up-regulated and down-regulated common DEGs between the Keto/Control and Keto/β-caro sets. MF, molecular function; BP, biological process; CC, cellular component; KEGG, KEGG pathway. The full list of enriched GO terms can be found in Table S12. Gene enrichment analysis was carried out with g:Profiler (https://biit.cs.ut.ee/gprofiler/).

Altogether, the transcriptomic data brought into light how the cells were adapting to the ketocarotenoids through modulating the expression of genes involved in the structure of the chromoplast, especially the membranes, the generation of precursor of metabolites and energy, the photosynthesis/chromorespiration and the redox control.

## DISCUSSION

### Cellular and metabolic adaptation to the heterologous formation of ketocarotenoids

The tomato plant, engineered to produce high levels of valuable ketocarotenoids, is an exemplar to study how the cell is able to respond and adapt to the biosynthesis of non-endogenous metabolites. In the present study, a series of adaptation mechanisms deployed by the cell were uncovered. Firstly, structural adaptations to increase the storage capacity for the sequestration of ketocarotenoids were discovered in the chloroplast and chromoplast membranes. More vesicles (membranous sacs) and a greater number of darker (electron dense) and larger plastoglobules were found in the EM of keto plastids (Figure 2a, 2c). A diversification of storage locations was also observed. While the predominant carotenoids of the control lines were only found in the membranes (membranous sacs and thylakoid remnants), the free and esterified ketocarotenoids were also detected in plastoglobules. The boost in membrane surface area is most likely a response to the increase in metabolite levels such as the ketocarotenoids, α-tocopherol and plastoquinone (Fig 1 and 2). Indeed, it was previously demonstrated that an increase in plastoquinone level led to enlarged plastoglobules (Szymanska and Kruk, 2010). Similarly, higher carotenoid levels resulted in altered membranous plastidial structures in many plant species (Enfissi et al., 2019, Kilcrease et al., 2013, Maass et al., 2009, Fu et al., 2012, Berry et al., 2019, Cao et al., 2012). Moreover, the plastid membrane composition was altered to presumably accommodate the ketocarotenoids. Indeed, the fatty acid composition of the main plastidial lipids (MGDG and DGDG) shifted from C36:3 to C36:6 (Figure 3c). The increase in the degree of unsaturation has been previously demonstrated as an adaptation mechanism to stresses such as cold temperature, whereby, the lipid transition temperature is lowered to maintain membrane fluidity, in effect stabilising the membranes (Leekumjorn et al., 2009, Guschina and Harwood, 2006, Menard et al., 2017, Wang et al., 2020). The esterification of (keto)carotenoids renders those metabolites more lipophilic, which favours their sequestration in membranous structures. In effect, this is also a mechanism to increase the carotenoid storage capacity of the plant cells (van Wijk and Kessler, 2017, Torres-Montilla and Rodriguez-Concepcion, 2021). The esterification state of the ketocarotenoids did not influence their localisation in the sub-compartments of the keto chromoplasts (Figure S5). Secondly, adaptation mechanisms involving the recruitment of promiscuous enzymes (esterification and catabolism of the ketocarotenoids, Figure 3a & 3b, Table S7) and/or enzymes whose genes are expressed under less regulatory control at a specific developmental stage, have also been witnessed. For example, the recruitment of the *Cyc-b*-Gal enzyme in the keto MG fruit instead of the favoured lycopene β-cyclase (*Lcy-b*, Table S7) at MG. Other examples of this adaptive mechanisms have been highlighted in the literature (Mortimer et al., 2016, Mortimer et al., 2017, Nogueira et al., 2013, Enfissi et al., 2017, Enfissi et al., 2019, Karniel et al., 2022). Thirdly, in order to cope with the synthesis of the ketocarotenoids, the core metabolic pathways across the cell have been mobilised for the generation of metabolite precursors and energy (Figure 4, 5 and 6). In particular the TCA cycle would seem to be providing reductants for mitochondrial respiration, fatty acid biosynthesis and possibly chromorespiration (Figure 4 and 6). The up-regulation of the expression of genes involved in the chromorespiration could also be linked to the increase of membranous surface in the chromoplasts, as chromorespiration is believed to take place in vesicles (Renato et al., 2014). Consequently, coordinated changes in all sub-compartments of the cell were observed. This implies that regulatory mechanisms were in place to orchestrate those global changes.

### Redox control as a global orchestrator of cell adaptation

The plant’s capacity to rapidly adapt to environmental cues relies principally on the quest for reduction-oxidation (redox) homeostasis at the cellular level. External stresses usually lead to a change in redox status, which is sensed, and redox regulation mechanisms are activated to restore homeostatic balance. The redox status plays a pivotal role in coordinating metabolic processes. Simultaneously, the redox status regulates itself through the metabolism with many metabolic pathways being targets of redox regulation. Consequently, the redox regulation of the metabolism results in modification of the redox status itself (Geigenberger and Fernie, 2014, Obata, 2015, Cejudo et al., 2019, Hernandez and Cejudo, 2021, Yokochi et al., 2021). In the present article, the plant cells were not reacting to external signals but to new intracellular cues in the form of non-endogenous metabolites. As a result of the high-level production of ketocarotenoids in the cell, a global response was initiated (Figure 4, 5 and 6). The intertwined relationship of redox control and metabolism suggests that redox control must have played a part in orchestrating those global changes observed and was in turn self-regulated by the redox changes occurring. The data in this article corroborated this hypothesis, as further described. For example, changes in redox related gene expression were observed (Figure 6, table S8). It is also known that NAD(P) and NAD(P)H couples are integrated in most of the core metabolic processes such as the glycolysis, the tricarboxylic acid (TCA) cycle, the respiratory electron transport, the photosynthesis and the oxidative pentose phosphate pathway (oxPPP) (Obata, 2015). All the mentioned pathways have been altered at a transcriptomic and/or metabolic level (Figure 4 & 6). Moreover, the transcript of NAD(P)H dehydrogenase subunit M (*Ndh-M*), which is part of a plastidial respiratory mechanism (Renato et al., 2015), was one of the most up-regulated genes (20-fold) in the ripe keto/control DEG. There was no difference in *Ndh-M* expression at the mature green stage between the keto and control fruit. However, while, in the control, its expression decreased (16-fold) through ripening, it remained constant in the keto fruit (Table S7). A similar trend was observed with other *Ndh* subunits (Table S8). Furthermore, the plastidial Ndh complex activity has been shown to impact central metabolism as well as antioxidant levels such as carotenoids, flavonoids and tocopherols (Nashilevitz et al., 2010). Hence, the changes observed in the ketocarotenoid tomato, namely the alteration of the core and specialised metabolism at metabolomic and transcriptomic level suggest that the Ndh complex could have played an important role in the response of the redox control. Additionally, the carotenoid pathway itself is directly connected to the redox status with some of its enzymes, phytoene desaturase (PDS) and zeta-carotene desaturase (ZDS), being part of the net electron transfer, while 1-deoxy-D-xylulose 5-phosphate reductoisomerase (DXR), lycopene β-cyclase (LCYB), β-carotene hydroxylase (CRTR-B) and Zeaxanthin epoxidase (ZEP) requiring NADPH or high PQ/PQH2 for their reactions (Figure 1b, Fanciullino et al., 2014, Nashilevitz et al., 2010). Therefore, alterations within the carotenoid pathway, which were triggered by the biosynthesis of ketocarotenoids, ought to have disrupted the redox homeostasis. The non-photochemical electron transport in the chromoplast, also called chromorespiration, is believed to involve, the Ndh complex, plastoquinone, the carotenoid enzyme PDS, PTOX, the cytochrome b6f and an ATP synthase (Renato et al., 2014, Grabsztunowicz et al., 2019). The plastoquinone pool is a redox sensor that regulates the redox status (Havaux, 2020). In particular, it has been shown in *Haematococcus*, that it is a redox sensor for carotenoid biosynthesis (Steinbrenner and Linden, 2003). Moreover, active exchange of plastoquinone between the thylakoid membrane and the plastoglobules is important for tight environmental control (Havaux, 2020). In stress conditions, a redistribution of plastoquinone towards the plastoglobules occurs, possibly due to the increased occupancy in the membrane which restrict plastoquinone diffusion (Kirchhoff, 2014). Here, our data showed an increased plastoquinone content in the keto tomato compared to the control line, which was previously hypothesised to be linked with the enlarged and darker plastoglobules observed in the keto EM (Figure 2). Moreover, hydrogen peroxide formation within the plastoquinone pool is thought to participate in retrograde signalling and in modulation of expression (Havaux, 2020). Hydrogen peroxide has also been suggested to play a key function in the control of the redox state of stromal enzymes (Cejudo et al., 2019) and signalling between cell sub-compartments (Buchanan and Balmer, 2005). The enriched GO term “response to hydrogen peroxide” in up-regulated DEGs of the keto tomato compared to the control (Table S6) reinforced the participation of the redox control in the cell response to ketocarotenoids. In addition, antioxidants, such as tocopherols, are considered information-rich redox buffers which react with a myriad of cellular components (Shao et al., 2008). The interaction between ROS and antioxidant is believed to act as integrator of signals derived from the environment and the metabolism (Foyer and Noctor, 2005, Shao et al., 2008). Therefore, the increase in tocopherol content in the keto tomato, had most likely an impact on redox signalling. Finally, the changes in TCA cycle metabolites studied (Figure 4) and the up-regulation of the expression of the plastidial malate dehydrogenase (Table S7) indicated a possible alteration of the malate valve to maintain redox homeostasis between sub-cellular compartments such as chloroplast and mitochondrion (Buchanan and Balmer, 2005, Selinski and Scheibe, 2019). Together these lines of evidence corroborate the involvement of redox control as part of an adaptation mechanism to the biosynthesis and storage of the ketocarotenoids.

### The impact of the ketocarotenoid production on photosynthesis and beyond

In comparison to previous ketocarotenoid forming lines such as the ZW tomato line (Enfissi et al., 2019), the inclusion of the *Solanum galapagense* lycopene β-cyclase (*Cyc-b*-Gal) restored β-carotene levels in the keto fruit studied (Figure 1, table S1 & S3). Interestingly, several differences were highlighted between the ZW line and the keto line (i) only the keto line could produce high levels of ketocarotenoids, due to the consistently maintained levels of the precursor β-carotene in the fruit; (ii) the ZW chloroplasts resembled chromoplasts due to the lack of thylakoid structures, while the keto line had wild type chloroplasts, with the addition of more vesicles; (iii) the ripening of the ZW tomatoes was delayed compared to that in the control tomatoes but no delay was observed in the keto tomatoes. Although the ZW tomatoes were producing 15 times less ketocarotenoids compared to keto tomatoes (Enfissi et al., 2019), the impact on the chloroplast sub-structures and ripening were a lot more pronounced. This hints at the importance of β-carotene in the global metabolism. Indeed some carotenoids, including β-carotene, are thought to be essential elements of photosynthesis. In the leaf, the Fv/Fm value (the maximum photochemical efficiency of PSII in the dark-adapted state) has been negatively impacted by the presence of the ketocarotenoids in the chloroplast membranes (Figure S3). This has been recorded in other ketocarotenoid engineered plants (Mortimer et al., 2017, Enfissi et al., 2019, Xu et al., 2020). Ketocarotenoids have been found to replace some or all carotenoids in the light harvesting complexes (Mortimer et al., 2017, Liguori et al., 2017), nevertheless, it has been proven that the photosynthetic system was still functional (Xu et al., 2020). The process of photosynthesis has shown a high adaptive capacity by being able to remain functional with only ketocarotenoids as accessory carotenoid pigments and, for instance, by compensating for the decrease in PSII/PSI antenna size in the keto engineered tobacco by a high PSII/PSI ratio (Xu et al., 2020). In the fruit, alteration of photosynthesis has also been demonstrated at the transcriptomic level with the up-regulation of photosynthetic genes (Figure 6) and the enriched GO terms linked to photosynthesis in the keto ripe tomato (Figure 5, table S6). In particular, the expression of the genes encoding the curvature thylakoid proteins has been up-regulated in the keto ripe fruit (Table S7). Those proteins have been suggested to play a role in the PSII repair cycle and in facilitating the connection between LHCII and PSI (Trotta et al., 2019). It is known that photosynthesis in the mature green tomato fruit is functional but is only fixing up to 20% of the carbon (Pesaresi et al., 2014, Simkin et al., 2020). Therefore, the fruits are still heavily dependent on the sugars provided by leaf photosynthesis (Lytovchenko et al., 2011). The photosynthesis in the keto leaf has been negatively impacted (Figure S4), similarly it could have been affected in the keto fruit but to a lesser extend as the fruit contained wild type level of β-carotene. Decreased sucrose content was found in the ripe keto tomato relative to the control (Figure 4, table S4). At a transcriptomic level, the photosynthesis genes seemed to have been first down-regulated in the MG keto tomato and then up-regulated in the ripe keto tomato (Figure 6, table S10). Rubisco (small unit), a main enzyme for the fixation of CO_2_ in the Calvin cycle, has been positively correlated (p-corr = 0.96) to a ketocarotenoid in a network analysis (Figure 7, Table S9). Its expression in the control fruit drastically decreased through ripening (35-fold), while in the keto fruit it remained constant (Table S7). It is possible that the photosynthesis in the fruit was compensating for the impacted photosynthesis in the struggling leaves. The photosynthesis compensation mechanism between plant compartments has already been shown. Indeed, the literature describes that any decrease in the rate of fruit photosynthesis can be compensated for by the up-regulation of leaf photosynthesis and an increased import of photoassimilates from the leaves (Nunes-Nesi et al., 2005, Araujo et al., 2011). The suggestion that the fruit photosynthesis could be up-regulated to boost the metabolism has implication for future genetic engineering of sink organs to enhance yield, fruit size and nutritional quality (Simkin et al., 2020). Moreover, the data described here suggest that other respiratory mechanisms have also been up-regulated to compensate for chemical energy synthesis such as chlororespiration and/or chromorespiration, which are both linked to, among others, NDH enzyme and plastoquinone levels (Figure 6). An extensive comparison of the ZW and keto tomato leaf and fruit would certainly shed more light on the plasticity of photosynthesis and other respiratory processes.

### Cell reprogramming or the quest for homeostasis

It is clear that in order to maintain cellular homeostasis holistic systems level changes arise in response to non-endogenous ketocarotenoid production. Deciphering the precise sequence of events will require further validation. However, Figure 9 has outlined the changes with direct and indirect effects, illustrating a potential cascade of events. For example, (i) the targeted carotenoid pathway is perturbed and transcriptional regulation is induced to regulate the carotenoid pathway flux (ii) related plastidial isoprenoids are effected potentially from ubiquitous precursor deviation. (iii) global changes across primary and intermediary metabolism arise, presumably to supply source precursors for complex energy consuming secondary metabolites, (iv) these metabolic changes, in the carotenoid pathway and beyond, are inevitably impacting the redox mechanism of the cell, which in turn is likely to drive secondary transcriptional and biochemical regulation and (v) the change in metabolite composition are directing organelle adaption in order to accommodate non-endogenous metabolites. Another feature that is evident arises from enzyme promiscuity, highlighted by the formation of carotenoid esters and new volatiles not reported in the volatome of tomato to date.

**Figure 9:**
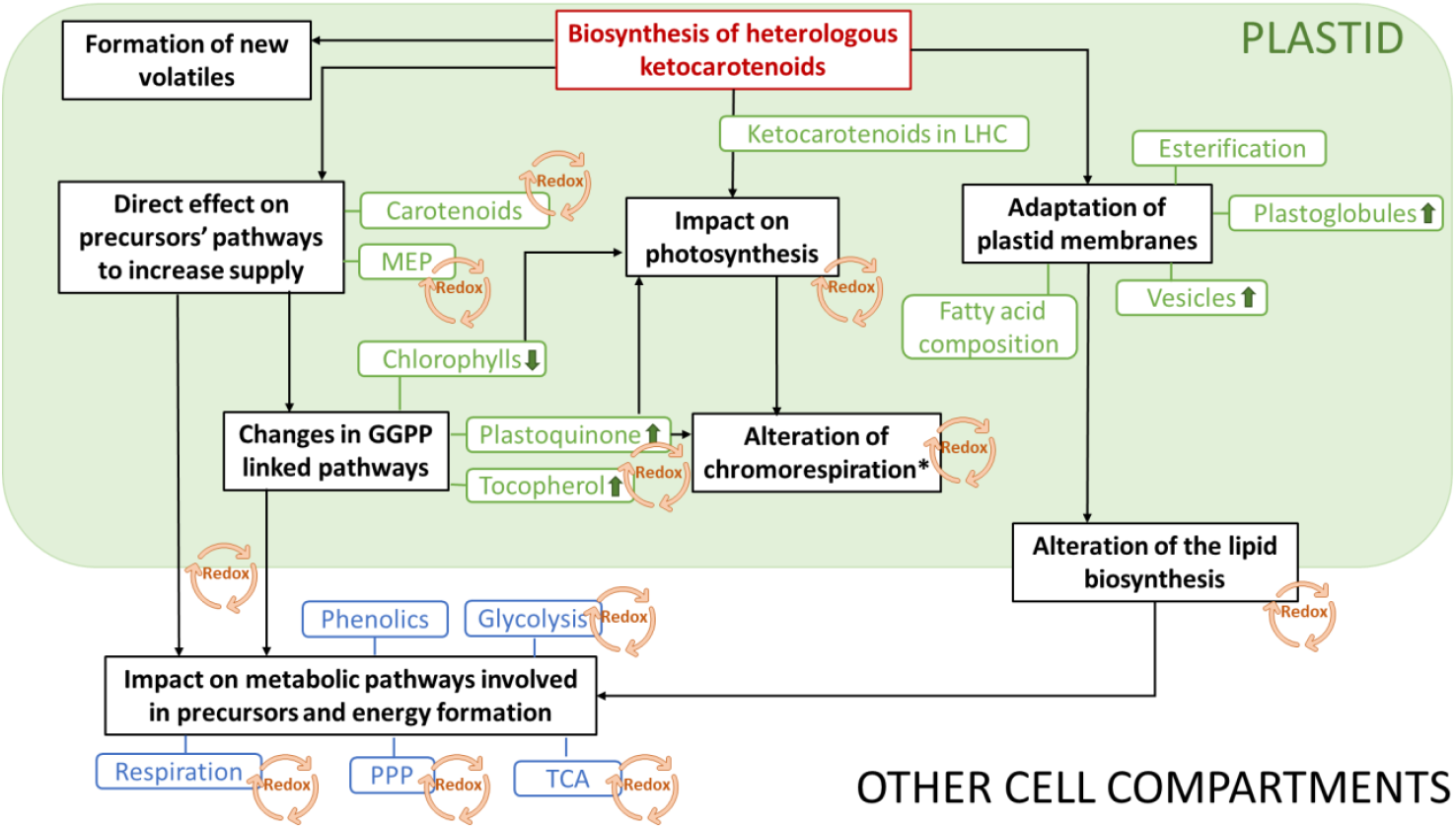
Transcriptomic, metabolic, and cellular reprogramming triggered by the production of heterologous ketocarotenoids. MEP, Methylerythritol 4-phosphate; GGPP, Geranylgeranyl pyrophosphate; PPP, Pentose phosphate pathway; TCA, Tricarboxylic acid cycle; LHC, Light-Harvesting Complex. Green arrows indicate an increased level. *, according to models proposed by Renato et al., 2014 and Grabsztunowicz et al., 2019. See main text for figure description.

In summary, our multi-omic approach has gone beyond the target pathway to show the complexity of metabolic reprogramming events that arise in an attempt to maintain homeostasis. Knowing the metabolic limits of the plant and how these limits could be extended, will aid in the rationale design of future engineering and precision breeding approaches promoting the bioeconomy. Transcription factors have been used and are routinely proposed as the key to coordinated gene expression, for optimal flux through a pathway (Zhang et al., 2015). This is in line with the standard representation of the central dogma and its proposed flow of information. However, in the present study it appears that production of non-endogenous metabolites are the initiators of global metabolic reprogramming, that employs multiple layers of cellular regulation, including transcription and organelle adaption. Interestingly, in animal systems cancerous tissues display metabolic reprogramming as a mechanism to attempt to maintain homeostasis, linked to the generation of energy. The metabolite changes observed in the ketocarotenoid producing lines are similar, whereby the nature of energy production, lipid and amino acid formation are perturbed, in addition to organelle biogenesis and macromolecular production occurring. Traditionally these changes in cancerous tissues have been attributed to induction of transcription factors (Ell and Kang, 2013, Sanda et al., 2012). However, alterations in specific metabolism/metabolite levels have more recently been postulated as the progenitors of transcriptional master regulators (Martin-Martin et al., 2018). The latter would appear to be the case in the present metabolic engineering study with non-endogenous compounds, initiating a series of metabolic reprogramming events. This regulatory process across kingdoms warrants further study and highlights the need to consolidate the role of metabolites as a component of the central dogma (Costa Dos Santos et al., 2021, de Lorenzo, 2014).

## Abbreviations

Dpa: days post anthesis
EM: electron micrograph
DEG: Differentially Expressed Genes
Keto: Ketocarotenoids
β-caro: β-carotene

## ACKNOWLEDGMENTS

This work was supported through the European Union Framework Program 7 DISCO project grant (613513) funded under the Knowledge-Based Bio-Economy program, Biotechnology and Biological Sciences Research Council OPTICAR Project (optimisation of tomato fruit carotenoid content for nutritional improvement and industrial exploitation; Project BB/P001742/1) and European Cooperation in Science and Technology (COST) Roxy-COST (Oxygen sensing a novel method for biology and technology of fruit quality; COST Action CA18210). We are grateful to IGATech and especially to Dr. F. Cattonaro and Dr. V. Vendramin for performing the RNAseq experiment and the cuffdiff. We are also thankful to Dr. Brown (University of Kent, UK) for the work provided in electron microscopy analysis of the ripe fruits and expertise.

## AUTHOR’S CONTRIBUTION

M.N., P.D.F, and E.M.A.E. designed the study and its conceptualisation, M.N., E.J.P, G.N.M, E.V. performed the research, M.N. analysed the data and M.N., P.D.F, and E.M.A.E wrote the manuscript with the input from all authors. PDF and E.M.A.E were responsible for the funding.

## METHODS

### Plant material and growth conditions

The ketocarotenoid, β-carotene and control tomato (*Solanum lycopersicum*) lines were generated in previous studies (Nogueira et al., 2017; Enfissi et al., 2019) where they are referred to as ZWRI, ZWØRI and ZWØRIØ, respectively. The tomato plants were greenhouse grown (25°C day/15°C night) at Royal Holloway University of London (England), with supplementary lighting (16 h day/8 h night).

### Metabolite extraction and analysis

Freeze-dried tomato material was homogenised to a fine powder with a tissue disruptor TissueRuptor (Qiagen). Extraction protocols varied for a minimum of three biological and three technical replicates and included one quality control and one extraction blank per randomised batch.

### Carotenoid, chlorophyll, and plastoquinone analyses

Tomato powder aliquots (15 mg) were extracted by the addition of chloroform and methanol (2:1). Samples were stored for 20 min on ice. Subsequently, water (1 vol.) was added. Samples were centrifuged for 5 min at top speed in a Heraeus Pico21 centrifuge (Thermo Scientific). The organic phase, containing the metabolites of interest, was placed in a fresh centrifuge tube and the aqueous phase was re-extracted with chloroform (x2 by volume). Organic phases were pooled and dried using the Genevac EZ.27 (SP scientific). Dried samples were stored at -20°C and resuspended in ethyl acetate or ethyl acetate/acetonitrile (3:7) prior to chromatographic analysis. Metabolites of interest were separated and identified by Ultra High Performance Liquid Chromatography with photo diode array detection (UPLC-PDA). An Acquity™ Ultra High Performance Liquid Chromatography UPLC System (Waters) was used with an Ethylene Bridged Hybrid (BEH C18) column (2.1 × 100 mm, 1.7 μm) with a BEH C18 VanGuard pre-column (2.1 × 50 mm, 1.7 μm). The mobile phase used was A: MeOH/H_2_O (50/50) and B: ACN (acetonitrile)/ethyl acetate (75:25). All solvents used were High Performance Liquid Chromatography HPLC grade. Several gradients were used depending on the metabolites. For the carotenoids, ketocarotenoids and chlorophylls (keto method), the gradient was 50% A: 50% B for 0.5 min and then stepped to 30% A: 70% B for 4.5 min, to 0% A: 100% B for 2 min, back to 30% A: 70% B for 1 min and to 50% A:50% B for the two last minutes. An extended version of keto method was used to analyse astaxanthin diesters (long keto method), with the 0% A: 100% B step lasting for 11 min instead of 2 min. For plastoquinone (caro method), the gradient was 30% A: 70% B for 0.5 min and then stepped to 0.1% A: 99.9% B for 5.5 min and then to 30% A: 70% B for the last 2 min. Column temperature was maintained at 30°C and the temperature of samples at 8°C. On-line scanning across the ultraviolet/visible range was performed in a continuous manner from 250 to 600 nm, using an extended wavelength photo diode array detector (Waters). Carotenoids, chlorophylls and plastoquinone were quantified using dose-response curves made using authentic standards. Content was expressed as μg/g DW.

### Fatty acids analysis

Fatty acids were extracted from 20 mg of freeze dried tomato powder. Transmethylation was performed as follow: Methanolic-HCl 1N (1 mL) was added to the material as well as the internal standard (Heptadecanoic acid, 100 μg) and incubated at 85°C for 3 h. Hexane (500 μL) and 0.8% potassium chloride (500 μL) were added to the mixture and vortexed twice for 10 sec. The mix was centrifuged for 5 min at 2,000 rpm. The upper phase was transferred into a fresh GC vial. Gas chromatography with flame-ionisation detector (GC-FID) analysis was performed on an Agilent 7890A with a DB-23 column. The gas chromatography oven was held for 2 min at 150°C before ramping at 10°C/min to 240°C. This final temperature was held for a further 9 min. Component identification was performed by comparison with fatty acid standards. Fatty acid quantification was achieved relative to the internal standard.

Fatty acid composition of digalactosyldiglycerol (DGDG) and monogalactosyldiglycerol (MGDG) were analysed using electrospray ionization tandem triple-quadrupole mass spectrometry (API 4000 QTRAP; Applied Biosystems; ESI-MS/MS). Total lipids were extracted from 20 mg of freeze dried tomato powder. Powder was resuspended using 4 ml chloroform/methanol 2:1 (v/v), vortexed and centrifuged 3 min at 3,500 rpm. KCl (1 ml, 1 M) was added and the solution was vortexed and centrifuged as previously. The upper phase was carefully discarded and 2 ml of milliQ water was added to the lower phase. The solution was again vortexed and centrifuged as previously. The lower phase was transferred in a new glass tube and dried down under nitrogen. Once dried the samples were resuspended in 150 μl of chloroform and stored under nitrogen at -20°C. An aliquot (25 μl) was combined with chloroform/methanol/300 mM ammonium acetate (300:665:3.5 v/v) to make a final volume of 1 mL. The lipid extracts were infused at 15 μL/min with an autosampler (HTS-xt PAL, CTC-PAL Analytics AG, Switzerland). The identification of the lipid species was performed using the scans neutral loss mode as follow: [M + NH4]+ in positive ion mode with NL 341.1 for DGDG and [M + NH4]+ in positive ion mode with NL 179.1 for MGDG. The scan speed was 1.5 sec/scan and 40 cycles were done per sample. The collision energies were +24 V for DGDG, and +21 V for MGDG. The MS scans were processed using the program Lipid View Software (AB-Sciex, Framingham, MA, USA) to determine the fatty composition of each MGDG and DGDG present in the samples.

### Primary/intermediary metabolism analysis

Tomato powder aliquots (10 mg) were extracted in 50% methanol (1 h, shaking, room temperature) followed by the addition of 1 volume of chloroform (500 μl). After, centrifugation, the non-polar extract was placed in a fresh centrifuge tube while the polar phase was re-extracted with 1 volume of chloroform. Polar (500 μl, methanolic epiphase) and non-polar metabolites (total: 1 ml, organic hypophase) were collected, after centrifugation. Saponified non-polar metabolites were obtained by modifying the protocol with the addition of 60% KOH (10X) in 50% methanol followed by an incubation at 42°C for 1 h in a thermomixer (300 rpm Eppendorf), instead of the 1 h rotation at RT. Polar extract (10 μl) and non-polar extracts (1 ml) were spiked with 10 μl of the internal standard solution (1 mg/ml of deuterated succinic acid in methanol and 1mg/ml of deuterated myristic acid in chloroform, respectively) and dried under vacuum (Genevac EZ.27). Dried extracts were derivatised to their methoxymated and silylated forms by adding 30 μl of methoxyamine hydrochloride (20 mg/mL, in pyridine; Sigma-Aldrich) followed by an incubation at 40°C for 1 h and then adding 70 μl of MSTFA (*N* methyl-*N*-trimethylsilyltrifluoroacetamide; Sigma-Aldrich) followed by another incubation at 40°C for 2 h. Gas chromatography-mass spectrometry analysis was performed on an Agilent 7890 (UK) gas chromatograph with a 5975C MSD. Samples (1 μl) were injected in a splitless mode at 290°C. Retention time locking to the internal standard was used. Metabolites were separated on a DB-5MS + DG 30 m (plus 10 m Duraguard) × 250 μ m × 0.25 μ m column (J&WScientific, Folsom, California, US). The GC oven was held for 1 min at 70 °C before ramping at 8 °C/min to 325 °C and held for 2 min. Helium was the carrier gas at a flowrate of 1 mL/min. The interface with the MS was set at 280 °C and MS performed in full scan mode using 70 eV EI + and scanned from 50 to 1000 m/z. A mixture of n-alkanes, ranging from 8 to 32 carbons, was used for retention index external calibration. Automated Mass Spectral Deconvolution and Identification System (AMDISv2.71, NIST) was used to create an in-house library based on authentic standards and NIST’11 MS library (National Institute of Standards and Technology, USA). Peak deconvolution was performed with AMDIS in batch mode for each sample set and peak identification conducted according to metabolomics reporting guidelines (Table S14). Peak areas were expressed relative to internal standard. Polar and non-polar datasets were combined; removing duplicates by selecting the maximal response recorded per each compounds present in both phases.

### Phenolics

Following the same extraction protocol as for primary metabolites, an aliquot of the polar extract was passed through nylon filters (0.45 μm), then transferred (100 μL) to glass inserts and spiked with an internal standard genistein (5 μL of 0.2 mg/mL). Analysis was performed with an Agilent 6560 Ion Mobility Q-TOF coupled to an Agilent 1290 Infinity II (Agilent Technologies,Inc.). The samples were separated with an YMC-UltraHTPro C18 column (100 × 2 mm i.d. 2 μm) using the LC–MS grade solvent A (water and 0.1 % formic acid) and solvent B (acetonitrile, 2.5% water and 0.1 % formic acid) at 0.2 mL/min. The gradient started at 95 % (A) for 0.5 min, followed by a linear decrease to 75 % (A) at 3 min, 70 % (A) at 6 min, 0% (A) at 6.5 min, which was held until 7.5 min before returning to initial conditions of 95 % (A) at 9.5 min. The column was then re-equilibrated for 1 min. The autosampler was kept at 8°C and the column at 30°C.The eluate was split for simultaneously analysis by DAD (scan mode 200−600 nm) and MS. Electrospray ionisation (ESI) was performed in negative mode with the capillary and nozzle set to 4,000 and -500 V, respectively. The nebulizer gas (nitrogen) was set at 35 psi, dry gas at 5 L/min and 325 °C and sheath gas at 12 L/min and 275 °C. The MS was run in full scan mode with a 100 to 1,700 m/z range at 0.9 spectra/second. Calibration to a reference solution was performed during each run. Molecular feature extraction was performed with Agilent Profinder (V10.0 SP1, Agilent Technologies, Inc.) with retention time tolerance of 0.3 min and mass tolerance of 10 ppm for peaks > 400 counts. Normalization was performed relative to the internal standard. Metabolites were identified by comparison of UV/VIS spectrum, mass spectrum and retention time to an in-house library of authentic standards. Peak areas were expressed relative to internal standard.

### Phytohormones

Isopropanol (Fisher Chemical) and water (2:1) with 0.1% HCl (500 μL, Fisher Chemical) was added to homogenised ripe tomato material (20 mg), samples were vortexed (10 sec) and rotated (1 h) at room temperature. Dichloromethane (Fisher Chemical) was added, samples vortexed (10 sec) and centrifuged (20,000 RCF, 5 min). Aliquots (100 μL) of the polar phase were transferred to glass inserts and dried under nitrogen. Samples were then resuspended in methanol (100 μL). Extracts were injected (1 μL) into a 1290 Infinity II UPLC (Agilent Technologies), coupled to a 6470 triple quadrupole (QQQ) MS (Agilent Technologies). The column, UPLC conditions, mobile phase and temperature were identical to the settings used for the flavonoid analysis. The solvent gradient was different. It started at 95 % (A) for 1 min, followed by a linear decrease to 75 % (A) at 3 min, 60 % (A) at 6 min, 2% (A) at 8 min, which was held until 9.5 min before returning to initial conditions of 95 % (A) at 10.5 min. The column was then re-equilibrated for 5 min. Samples were simultaneously analysed in ESI+ and ESI-modes, with capillary voltage set to 3,500 V and 4000 V, respectively, and nozzle voltage of 2,000 V and -500 V, respectively. The N2 drying gas flow rate was 11 L min-1 and the cell accelerator voltage was 5 V. A dynamic multiple reaction monitoring (MRM) method was used to selectively analyse the phytohormones of interest. Details for each phytohormone are given as retention time (min), scan mode, collision energy (V), fragmentor (V), precursor ion (m/z) and product ion (m/z): Zeatin: 2.4 min, Negative, 19 V, 80 V, 218.10 m/z, 134.0 m/z; Gibberellic acid 3 (GA3): 3.88 min, Negative, 34 V, 110 V, 345.13 m/z, 143.1 m/z; Indole-3-acetic-acid (IAA): 4.78 min, Negative, 10 V, 90 V, 174.05 m/z, 130.1 m/z; Salicylic acid (SA): 4.85 min, Negative, 18 V, 90 V, 137.02 m/z, 93.1 m/z; Abscisic acid (ABA): 5.15 min, Negative, 10 V, 90 V, 263.13 m/z, 153.1 m/z; Jasmonic acid (JA): 6.16 min, Negative, 10 V, 90 V, 209.12 m/z, 59.1 m/z; Gibberellic acid 4 (GA4): 7.06 min, Negative, 34 V, 80 V, 331.5 m/z, 213.1 m/z; Jasmonic acid isoleucine (JA-Ile): 7.20 min, Negative, 22 V, 80 V, 322.20 m/z, 130.1 m/z; Methyl jasmonic acid (MeJA): 7.70 min, Positive, 10V, 90 V, 225.15 m/z, 151.1 m/z. Ion chromatograms were extracted and integrated for each phytohormone product ion in MassHunter Qualitative Analysis 10.0 (Agilent Technologies). Peak areas were converted to concentrations (ng/g DW) via external standard calibration curves produced with authentic standards.

### Volatiles

Tomato fruits frozen in liquid nitrogen were ground to a fine powder in mortar and pestle and 2 g transferred to a pre-cooled (on ice) 20 mL opaque glass vial. Samples of air taken during sample preparation were analysed as blanks. Samples were stored at -20 °C for 1 h prior to analysis. Vials were loaded onto a GC-MS system with SPME capable autosampler. Samples were incubated for 30 min at 60 °C whilst agitating at 300 rpm then headspace analytes extracted onto 1 cm, 23 ga, 50/ 30 μm Car/DVB/PDSM fibres for 20 min under same conditions. Following this, analytes were desorbed from the fibre into the GC inlet, equipped with 0.75 mm internal diameter liner, at 250 °C for 3 min. A 5 min post-desorption bake-out at 250 °C was used to re-condition fibres. Analytes were resolved on a DB-5MS 30 m x 250 μm x 0.25 μm column with a temperature ramp as follows: 40 °C for 1 min then ramped to 250 °C at 5 °C/ min with a 2 min hold at both 120 °C and 250 °C, followed by a ramp to 300 °C at 6 °C/min and held for a further 5 min. Helium was the carrier gas at 1 mL/ min. Transfer line to the quadrupole MS was kept at 250 °C and spectra measured following EI+ at 70eV and operating in full scan mode from 33 – 600 m/z. Chromatographic data were processed via AMDIS software (NIST) and compounds identified against an in-house library of authentic standards and NIST ‘11 database (spectral match > 800) using Kovat’s retention indices generated following analysis of a C_5_-C_23_ alkane series under same conditions.

Detection of oxo-β-ionone was confirmed by another analytical method. Seventy gram of frozen fruit tissue was ground in liquid N_2_ into a fine powder. The volatile components were extracted with 100 ml of methyl tert-butyl ether (MTBE) by vigorous shaking on a shaker apparatus overnight. For quantitation, 10 μg of isobutyl benzene was added. The upper MTBE layer was separated, dried with sodium sulfate and concentrated to 1 ml. The samples were kept at 4°C until analysis. A 1 μl aliquot of the MTBE extract was injected into an Agilent GC 6890 system, coupled to quadrupole mass spectrometer detector 5973N (CA, USA). The instrument was equipped with Rxi-5sil MS column (30 m length × 0.25 mm i.d., 0.25 μm film thickness, stationary phase 95% dimethyl-5% diphenyl polysiloxane). Helium (11.6 psi) was used as a carrier gas with splitless injection. The injector temperature was 250 °C, and the detector temperature was 280 °C. The following conditions were used: initial temperature 50 °C for 1 min, followed by a ramp of 5 °C/min to 180 °C, and 10 °C/min up to 280 °C (5 min). A quadrupole mass detector with electron ionization at 70 eV was used to acquire the MS data in the range of 41 to 350 m/z. A mixture of straight-chain alkanes (C7-C23) was injected into the column under the above-mentioned conditions for determination of retention indices. The identification of the volatiles was assigned by comparison of their retention indices with those of literature and by comparison of spectral data with standard or with the Nist 98 and QuadLib 2205 GC-MS libraries. Component amount in each sample was calculated as (peak area x internal standard response factor) divided by (response factor x internal standard peak area). Oxo-β-ionone was quantified in the keto line (14 ± 2 ng compound/gr tissue/vial) and not detected in the control line.

### Semi-synthesis and identification of carotenoid-derived volatiles

Ketolated derivates of β-ionone and β-cyclocitral were generated via bromination with DBDMH (1,3-dibromo-5,5-dimethylhydrantoin) and subsequent oxygenation. A 1:1.1:1.5 molar ratio (substrate:DBDMH:NaOH) reaction was stirred at RT (∼22 °C) for 5 h whilst protected from light (黎 成 勇 *et al*., 2015). Products were extracted into MtBE. Hydroxylated derivatives were generated following reduction of the ketolated derivatives via sodium borohydride. Products were confirmed through EI+ GC-MS via liquid injection. Dried residue (200 μg) of authentic standards (lycopene, β-carotene, phoenicoxanthin, canthaxanthin, zeaxanthin) in 20 mL headspace were analysed under same condition as tomato fruit samples and manual comparative interpretation of mass spectra undertaken to enhance annotation of carotenoid cleavage products.

Identification features used for all compounds analysed are given in Table S14.

### Subchromoplast fractionation on a sucrose gradient

Pericarp, from 10 tomatoes at breaker + 5 days ripening stage, was cut into 1 cm^2^ pieces (80 to 150 g) and stored at 4°C overnight. From here, work was carried out at 4°C. Tissue was immersed in extraction buffer (0.4 M sucrose, 50 mM Tris, pH 7.8, 1 mM EDTA, and 1 mM DTT) and homogenised for 2 × 3 s in a Waring blender. The homogenate was then filtered through four layers of muslin. Subsequently, extraction buffer was added to the filtrate in centrifuge tube (500 ml). Tubes were centrifuged for 10 min at 5,000 g in a Sorvall RC5C centrifuge (Thermo Scientific) with a GSA-3 rotor. The supernatant was discarded. The pellet was resuspended in extraction buffer and transferred into smaller centrifuge tubes (50 ml). The tubes were centrifuged for 10 min at 9,000 g with a GSA-5 rotor. The supernatant was removed. Pellets were resuspended in 45% sucrose buffer (45% [w/v] sucrose, 50 mM Tricine, 2 mM EDTA, 2 mM DTT, and 5 mM sodium bisulphite, pH 7.9; 3 ml). The chromoplasts were physically broken using a handheld potter homogenizer (10 times). The solution was then resuspended in 45% sucrose buffer (5 ml). A total volume of 8 ml was placed in a 38.5-mL Ultra-Clear centrifuge tube (Beckmann Coulter). Subsequently, other layers of discontinuous sucrose gradient were overlaid, consisting of 38% sucrose buffer (6 ml), then 20% sucrose buffer (6 ml), 15% sucrose buffer (4 ml), and 5% sucrose buffer (8 ml). Gradients were centrifuged at 4°C for 17 h at 100,000 g using an L7 ultracentrifuge with an SW28 swing out rotor (Beckman Coulter). Fractions (F, 1 ml) were collected, from the top of the gradients using a Minipuls 3 peristaltic pump and FC203B fraction collector (Gilson). The identification of the different subchromoplast elements (plastoglobules and membranes) was previously demonstrated by western blots in Nogueira et al. (2013).

### Determination of in vivo chlorophyll fluorescence

In vivo chlorophyll fluorescence was measured using a pocket PEA chlorophyll fluorimeter (Hansatech Instruments, King’s Lynn, UK). Minimal fluorescence (Fo) was determined by applying a weak modulated light (0.4 μmol photons m−2.s−1) and maximal fluorescence (Fm) was induced by a short pulse (0.8 s) of saturating light (9000 μmol photons m−2.s−1). Maximal photochemical efficiency of PSII (Fv/Fm) was calculated as Fv/Fm = (Fm-F0)/Fm. Records were carried out on attached leaves in a dark-adapted state.

### Transmission electron microscopy

Tomato fruit was cut into 3 mm cubes using a razor blade on a tile, they were placed in a vial containing (∼3 ml) room temperature (RT) fixative (2.5% glutaraldehyde in 0.01M phosphate buffered saline (PBS) at pH 7.2) and placed under vacuum for 20 seconds to aid infiltration. Samples were placed on a rotator for ∼5 hours before going in the fridge (4°C) overnight. Tissue was washed in PBS 2 × 20 mins and then postfixed in 1% osmium tetroxide in PBS for 1hr at RT. Tissue was then dehydrated in increasing concentrations of acetone as follows: 10%, 30%, 50%, 70%, and 90% and 3 × 100% (minimum 1hr for each wash). Tissue was then infiltrated in increasing concentrations of resin diluted with acetone and placed under vacuum for 20 seconds as follows: 30%, 70% and 2 × 100% Spurr resin (minimum 1.5 hrs for each wash). Spurr medium ERL4221D (TAAB Laboratories Equipment Ltd.) was used and freshly mixed on the day. Tissue pieces were then placed in labelled capsules containing fresh resin and polymerized in the oven (60°C) for 16 hrs. Polymerized blocks were sectioned at 70nm on a Leica Ultracut UC7, sections were collected on 200 mesh copper grids coated with carbon-formvar. Sections were counterstained with 2.5% uranyl acetate in distilled water for 20 mins, washed with distilled water, stained with Reynolds lead citrate for 3 mins and then washed with distilled water again. Regions of interest were viewed in a JEOL JEM-2100 Plus Transmission Electron Microscope with an accelerating voltage of 200kV. Images were recorded with a Gatan OneView IS camera. The images shown in Figure 2 are representative of three biological replicates for each line, from which 12-20 regions of interest were imaged per biological replicate. Plastid measurements were determined using ImageJ software. In particular, the colour intensity of the plastoglobules (grey ratio) was calculating by measuring the colour intensity within several plastoglobules at three locations within the plastoglobules of an image and normalising this value to the colour intensity of the background (stroma) of this image (9 positions per image).

### RNA-seq

RNeasy Plant Mini kit (Qiagen) was used to extract total RNA from 39 and 49 dpa tomato fruit following manufacturer’s instructions. Single-read mRNA sequencing (1×100bp) was carried out with Illumina HiSeq2500. TruSeq Stranded mRNA Sample Prep kit (Illumina, San Diego, CA) has been used for library preparation following the manufacturer’s instructions, starting with 1-2ug of good quality RNA (R.I.N. >7) as input. The poly-A mRNA was fragmented 3 minutes at 94°C and every purification step has been performed by using 1X Agencourt AMPure XP beads. Both RNA samples and final libraries were quantified by using the Qubit 2.0 Fluorometer (Invitrogen, Carlsbad, CA) and quality tested by Agilent 2100 Bioanalyzer RNA Nano assay (Agilent technologies, Santa Clara, CA). Libraries were then processed with Illumina cBot for cluster generation on the flowcell, following the manufacturer’s instructions and sequenced on single-end mode at the multiplexing level requested on HiSeq2500 (Illumina, San Diego, CA). The CASAVA 1.8.2 version of the Illumina pipeline was used to processed raw data for both format conversion and de-multiplexing. RNA-Seq standard bioinformatics analysis were performed including base calling and demultiplexing, trimming (with ERNE1 and Cutadapt2), alignments with TopHat2, transcripts count (Cufflinks3), quality control using RSeqQC4 package, pair-wise differential expression analysis (Cuffdiff5). Mapping of reads to the Solanum lycopersicum SL3 (NCBI_SL3.0_protein) genome and gene ontology annotation was achieved with OmicsBox software as well as the Gene Ontology enrichment analyses (Fisher’s Exact Test, FDR ≤0.05) comparing two sets of DEGs. The Gene Ontology enrichment analysis of a single DEG set was completed on g:Profiler web server.

### Correlation network

Computations of the correlations were conducted in XLSTAT. Pearson correlation analysis was employed to compute all pairwise correlations between log2 of fold change metabolite levels and log2 of fold change expression levels for significantly different metabolites and DEGs of the ketocarotenoid line compared to the control (p-value < 0.5). Cytoscape was used to generate graphical output of network.

### Statistical analysis

Analyses were performed on datasets comprising three to 6 biological replicates and at least three technical replicates. N represented the number of biological replicates i.e. the number of plants studied per line. Analyses were made upon three pooled fruits per biological replicates. The technical replicates consisted in multiple extractions of each biological replicate. Absolute quantification was achieved by using dose response curves of authentic standards. When standards were not commercially available, relative quantification was performed. Statistical analyses were carried out using SPPS 25 software (IBM) and XLSTAT (Addinsoft). Comparison of two and more than two lines were subjected to Independent Samples T-test and

One-Way Anova statistical analysis, respectively. Equality of variances was tested with a Levene’s test. If the homogeneity of variance was assumed the p-value selected was, for T-test, the one for “equal variance assumed” and for ANOVA, the one of the Tukey post Hoc test. However, if the assumption of the homogeneity of variance was violated, then the p-value corresponding to “equal variance not assumed” was chosen for the T-test and for the ANOVA, the one corresponding to the Games-Howell post Hoc. Means and standard deviation were calculated in Excel as well as Pearson correlation (p-corr, p-value). SIMCA 17 (Sartorius) was used to carry out and display clusters derived from univariate scaling applied data for principal component analysis (PCA) and to perform “bottom up” hierarchal clustering. P-value of statistical tests performed are described in Table S13 if not already presented as supplemental data. Randomization technique was used whenever possible, such as running GC-MS and LC-MS samples in a random order. Microsoft Excel software was used to create randomized sequences.

## DATA AVAIILABILITY

Raw RNA-seq data have been deposited at the European Nucleotide Archive (Project accession PRJEB57363) and are publicly available as of the date of publication.

