## Supplemental information_1 for "Ketocarotenoid production in tomato triggers metabolic reprogramming and cellular adaptation: The quest for homeostasis?"

Nogueira et al.,

**<u>SUPPLEMENTAL INFORMATION</u>**

| | µg/g DW | 25dpa | | | 39dpa | | | 43dpa | | | 49dpa | | | 66dpa | | |
|---|---|---|---|---|---|---|---|---|---|---|---|---|---|---|---|---|
| **CONTROL** | **Neo/viola** | 72.9 | ± | *10.1* | 63.7 | ± | *4.1* | 62.5 | ± | *5.8* | 0.0 | ± | *0.0* | 55.3 | ± | *1.1* |
| | **Lutein** | 169.7 | ± | *15.2* | 172.2 | ± | *17.0* | 176.1 | ± | *2.4* | 161.8 | ± | *7.7* | 148.9 | ± | *1.0* |
| | **cis-Lycopene** | 0.0 | ± | *0.0* | 0.0 | ± | *0.0* | 0.0 | ± | *0.0* | 76.5 | ± | *2.2* | 55.7 | ± | *2.5* |
| | **Lycopene** | 0.0 | ± | *0.0* | 0.0 | ± | *0.0* | 35.1 | ± | *31.2* | 1903.3 | ± | *170.4* | 2644.0 | ± | *85.3* |
| | **γ-Carotene** | 0.0 | ± | *0.0* | 0.0 | ± | *0.0* | 0.0 | ± | *0.0* | 71.0 | ± | *3.5* | 92.6 | ± | *2.7* |
| | **β-Carotene** | 62.4 | ± | *1.4* | 64.1 | ± | *3.4* | 89.4 | ± | *12.9* | 164.3 | ± | *6.7* | 168.8 | ± | *8.2* |
| | **Total carotenoids** | 305.0 | ± | 26.6 | 300.0 | ± | 22.3 | 363.2 | ± | 35.4 | 2376.9 | ± | 172.9 | 3165.3 | ± | 76.0 |
| **β-CAROTENE LINE** | **Neo/viola** | 84.5 | ± | *5.9* | 74.9 | ± | *3.8* | 65.8 | ± | *4.0* | 56.2 | ± | *3.2* | 59.0 | ± | *4.2* |
| | **Lutein** | 180.0 | ± | *9.2* | 168.3 | ± | *10.9* | 154.2 | ± | *7.3* | 135.3 | ± | *8.6* | 131.0 | ± | *3.2* |
| | **Lycopene** | 0.0 | ± | *0.0* | 0.0 | ± | *0.0* | 0.0 | ± | *0.0* | 59.6 | ± | *12.8* | 50.3 | ± | *3.8* |
| | **α-carotene** | 0.0 | ± | *0.0* | 0.0 | ± | *0.0* | 0.0 | ± | *0.0* | 71.8 | ± | *6.3* | 105.8 | ± | *10.4* |
| | **γ-Carotene** | 0.0 | ± | *0.0* | 0.0 | ± | *0.0* | 0.0 | ± | *0.0* | 94.4 | ± | *19.8* | 71.4 | ± | *11.8* |
| | **β-Carotene** | 66.0 | ± | *1.2* | 62.8 | ± | *0.9* | 151.0 | ± | *73.7* | 954.8 | ± | *200.6* | 1014.9 | ± | *312.9* |
| | **Total carotenoids** | 330.4 | ± | 16.2 | 306.1 | ± | 14.6 | 370.9 | ± | 78.5 | 1372.0 | ± | 215.9 | 1432.4 | ± | 329.0 |
| **KETOCAROTENOID LINE** | **Astaxanthin** | 53.1 | ± | *8.4* | 40.8 | ± | *9.6* | 25.5 | ± | *2.9* | 37.2 | ± | *6.5* | 76.3 | ± | *14.9* |
| | **Phoenicoxanthin** | 26.1 | ± | *10.2* | 7.8 | ± | *11.5* | 49.1 | ± | *19.9* | 209.9 | ± | *16.2* | 518.9 | ± | *84.9* |
| | **Canthaxanthin** | 22.0 | ± | *14.3* | 6.8 | ± | *9.3* | 132.9 | ± | *76.3* | 558.2 | ± | *111.5* | 801.6 | ± | *151.5* |
| | **3'-OH-Echinenone** | 13.4 | ± | *1.0* | 12.4 | ± | *0.6* | 21.5 | ± | *6.6* | 78.5 | ± | *17.9* | 131.5 | ± | *33.9* |
| | **Echinenone** | 14.1 | ± | *1.0* | 14.1 | ± | *0.4* | 19.8 | ± | *5.3* | 51.9 | ± | *5.9* | 92.5 | ± | *9.8* |
| | **Phoenicoxanthin-C14:0** | 1.4 | ± | *0.0* | 1.5 | ± | *0.0* | 7.5 | ± | *6.0* | 128.2 | ± | *21.7* | 335.0 | ± | *187.6* |
| | **Adonixanthin-C14:1** | 0.0 | ± | *0.0* | 0.0 | ± | *0.0* | 11.5 | ± | *4.5* | 44.8 | ± | *7.2* | 153.0 | ± | *63.9* |
| | **Phoenicoxanthin-C16:0** | 2.2 | ± | *0.1* | 2.2 | ± | *0.2* | 21.2 | ± | *13.9* | 117.8 | ± | *22.5* | 203.3 | ± | *79.0* |
| | **Adonixanthin-C16:1** | 7.6 | ± | *0.4* | 8.4 | ± | *0.2* | 15.6 | ± | *3.1* | 44.1 | ± | *7.3* | 86.3 | ± | *22.9* |
| | **Adonixanthin epoxide** | 62.8 | ± | *4.7* | 63.6 | ± | *2.5* | 56.1 | ± | *3.0* | 0.0 | ± | *0.0* | 70.6 | ± | *0.8* |
| | **Total ketocarotenoids** | 202.6 | ± | 21.6 | 157.5 | ± | 21.7 | 360.9 | ± | 134.6 | 1270.5 | ± | 197.0 | 2469.0 | ± | 572.0 |
| | **Lutein** | 130.1 | ± | *1.7* | 126.6 | ± | *2.7* | 123.7 | ± | *2.0* | 0.0 | ± | *0.0* | 0.0 | ± | *0.0* |
| | **β-Carotene** | 55.4 | ± | *1.0* | 62.5 | ± | *3.5* | 73.9 | ± | *12.1* | 169.3 | ± | *25.9* | 240.3 | ± | *75.5* |
| | **Total carotenoids** | 185.4 | ± | 2.4 | 189.1 | ± | 5.3 | 197.6 | ± | 12.1 | 169.3 | ± | 25.9 | 240.3 | ± | 75.5 |

| | | | | | | |
|---|---|---|---|---|---|---|
| | **Total caro + keto** | 388.0 ± 23.4 | 346.7 ± 26.4 | 558.5 ± 145.4 | 1439.9 ± 173.2 | 2709.3 ± 540.6 |
| | **% Phoenicoxanthin esterified/Total phoenico** | 12 | 32 | 37 | 54 | 51 |

**Table S1: Carotenoid quantification in fruit over 5 ripening stages**

Carotenoid content is presented as mean in µg/g dry weight ± SD. Three representative fruits from three plants were used. Fruits were pooled and three determinations were made per sample. Results of statistical analysis are compiled in Table S13.

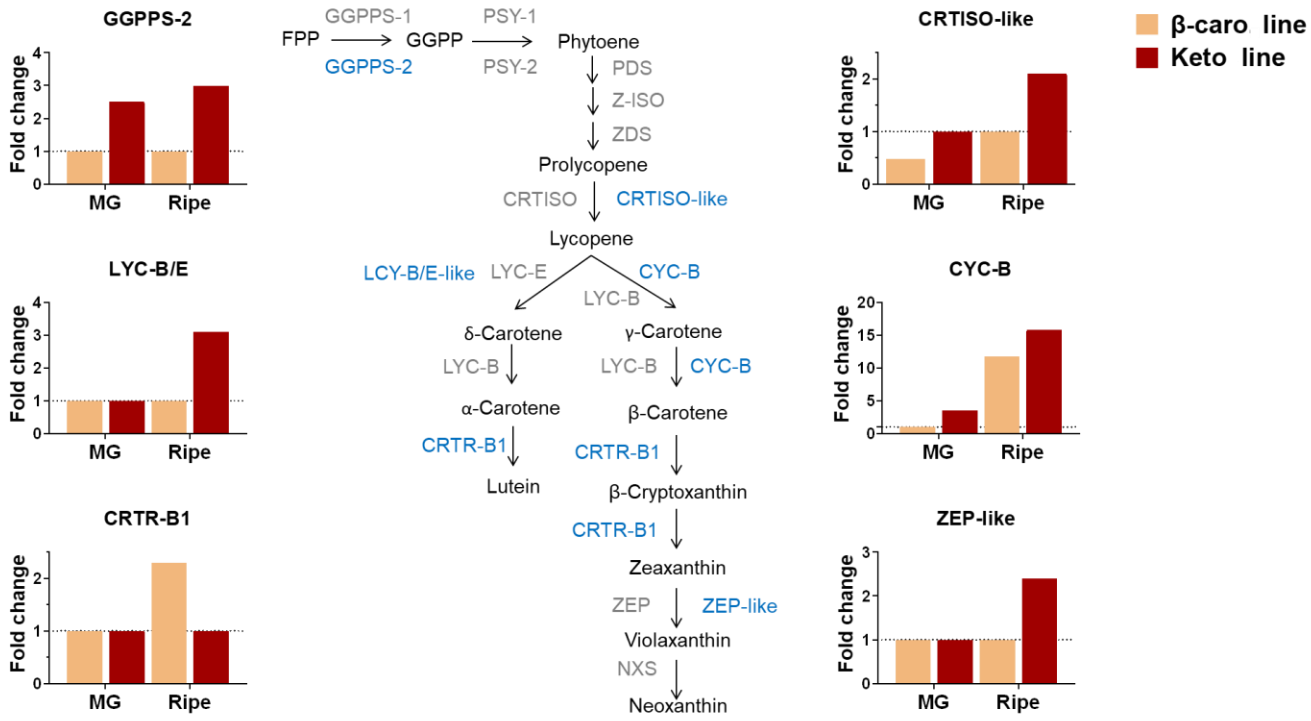


**Figure S1: Differential expression of the carotenogenic genes in the ketocarotenoid MG and ripe fruits compared to their respective controls**

A selection of the RNAseq data is displayed over the biosynthetic carotenoid pathway. Molecules are in black and enzymes are in capital letters, in grey and blue font. The latter depicts a significant change of expression level for the gene ($P < 0.05$), which encodes the enzyme described on the pathway. Data represent the fold change of expression level of a gene in the keto fruit compared to the control fruit (dashed line =1) for two ripening stages (Mature green (MG) and Ripe). FPP, farnesyl diphosphate; GGPP, geranylgeranyl diphosphate; GGPPS-1 and -2, geranyl diphosphate synthase; PSY-1, fruit specific phytoene synthase-1, PSY-2, phytoene synthase-2; PDS, phytoene desaturase; ZDS, ζ-carotene desaturase, CRTISO, carotene isomerase; CRTISO-like, carotene isomerase like (Solyc05g010180); LCY-E, ε–lycopene cyclase; LCY-B, β-lycopene cyclase; LCY-B/E, β/ε–lycopene cyclase (Solyc01g102950) ; β-CYC-B, fruit specific β-lycopene cyclase; CRTR-B1, carotene β-hydroxylase 1; ZEP, zeaxanthin epoxidase; ZEP-like, zeaxanthin epoxidase like (Solyc01g009080); NXS, neoxanthin synthase.

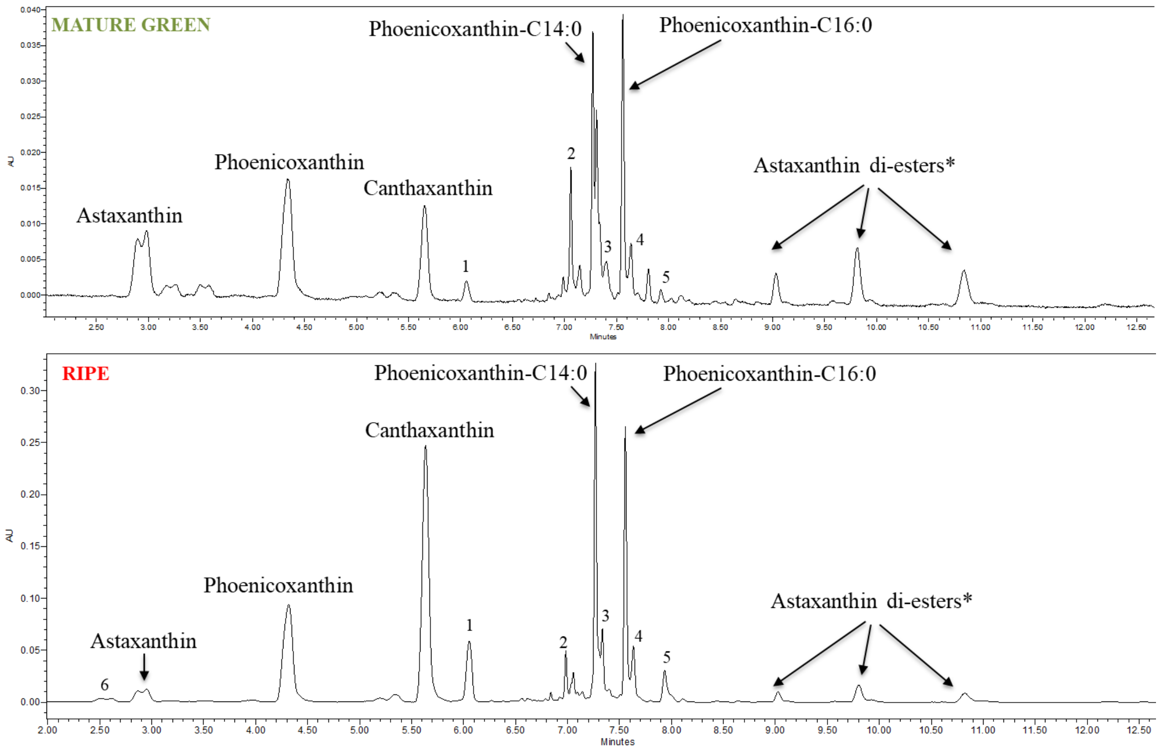


**Figure S2: Ketocarotenoid chromatographic profiles of the ketocarotenoid fruit at mature green and ripe stages**

Chromatograms were obtained by UPLC analysis (see methods) and recorded at 470 nm. 1, 3'OH-Echineone; 2, Echinenone; 3, Adonixanthin-C14:1; 4, Adonixanthin-C16:1; 5, β-Carotene; 6, Adonixanthin epoxide; *, putative

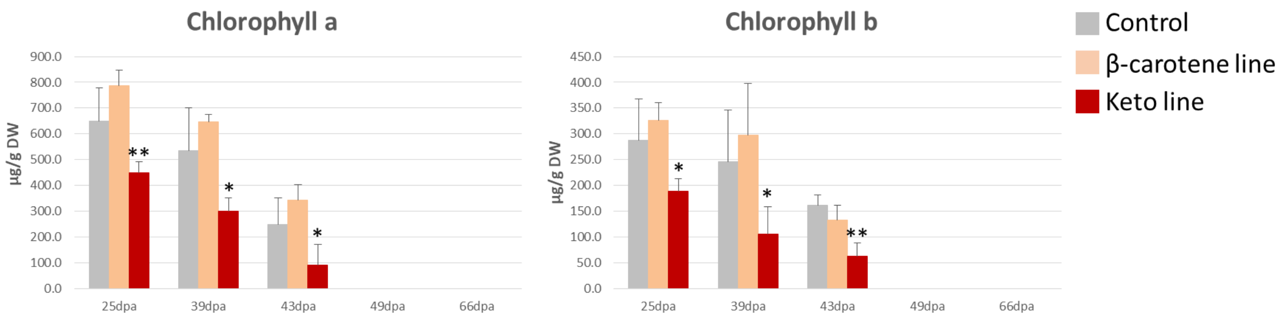


**Figure S3: Chlorophyll content in fruit at different developmental and ripening stages**

Chlorophyll a and b contents are presented as mean in µg/g dry weight ± SD. Three representative fruits from three plants were used. Fruits were pooled and three determinations were made per sample. Asterisks indicate significant differences compared to the control line (ANOVA analysis). $P < 0.05$, $P < 0.01$, and $P < 0.001$ are designated by *, **, and ***, respectively. P-values are listed in Table S13. Significant differences in the keto line compared to the β-Caro line were as follow: Chlorophyll a: 25 dpa: **; 39 dpa: **; 43 dpa: *; Chlorophyll b: 25 dpa: *; 39 dpa: ns; 43 dpa: *.

A

| mg/g DW | Control | β-caro. line | Keto. line | Keto. line / Control |
|---|---|---|---|---|
| Neoxanthin | 1.3 ± 0.04 | 1.3 ± 0.03 | **0.5 ± 0.00** | 0.4 |
| Violaxanthin | 1.6 ± 0.11 | 1.6 ± 0.04 | **0.5 ± 0.00** | 0.3 |
| Lutein | 4.4 ± 0.17 | 4.4 ± 0.12 | **1.8 ± 0.03** | 0.4 |
| β-Carotene | 1.3 ± 0.07 | 1.3 ± 0.05 | **0.5 ± 0.00** | 0.4 |
| Astaxanthin | ND | ND | **0.2 ± 0.05** | ∞ |
| Adonixanthin | ND | ND | **0.1 ± 0.00** | ∞ |
| Phoenicoxanthin | ND | ND | **0.7 ± 0.04** | ∞ |
| Canthaxanthin | ND | ND | **1.5 ± 0.09** | ∞ |
| 3'OH-Echinenone | ND | ND | **0.2 ± 0.00** | ∞ |
| Echinenone | ND | ND | **0.2 ± 0.01** | ∞ |
| Adonixanthin epoxide | ND | ND | **0.7 ± 0.05** | ∞ |
| Zeaxanthin | ND | ND | **1.5 ± 0.02** | ∞ |
| Chlorophyll b | 20.6 ± 0.84 | 19.6 ± 0.85 | **14.9 ± 0.17** | 0.7 |
| Chlorophyll a | 41.5 ± 2.25 | 40.9 ± 1.95 | **30.7 ± 0.34** | 0.7 |

B

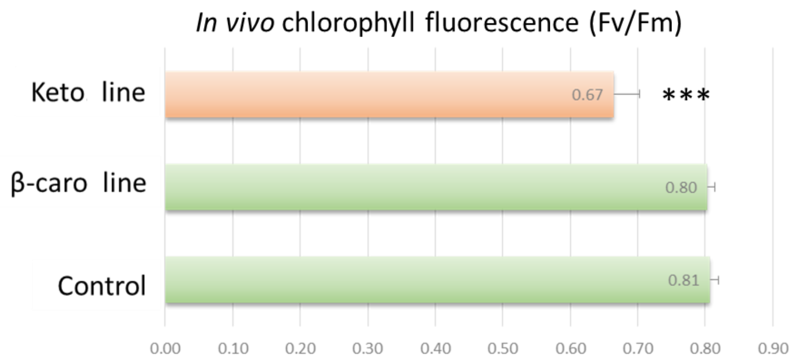


**Figure S4: Quantification of carotenoids and chlorophylls in leaves and measurement of *in vivo* chlorophyll fluorescence (Fv/Fm)**

(A) Carotenoid and chlorophyll contents are presented as mean in mg/g dry weight ± SD. Three representative leaves from three plants were used. Leaves were pooled and three determinations were made. Significant differences between the keto line compared to the other lines are indicated in bold (p-value < 0.001). ND, non-detected; ∞, only present in the keto line. (B) Measurement of *in vivo* chlorophyll fluorescence was done on six representative attached leaves from five plants for each genotype. Fv/Fm represents the maximum photochemical efficiency of PSII in the dark-adapted state and was calculated from the Kautsky Induction curve (see methods). Asterisks indicate significant differences compared to the control line (ANOVA analysis). P < 0.05, P < 0.01, and P < 0.001 are designated by *, **, and ***, respectively. P-values are listed in Table S13.

A

| MG | % of plastids containing X membranous sacs | | |
|---|---|---|---|
| X= | ≤10 | >10 | total % |
| **Control** | 32 | 6 | 37 |
| **β-caro. line** | 22 | 6 | 28 |
| **Keto. line** | 19 | **47**** | **66*** |

| RIPE | % of plastids with containing X membranous sacs | | |
|---|---|---|---|
| X= | ≤10 | >10 | total % |
| **Control** | 0 | 0 | 0 |
| **β-caro. line** | 0 | 0 | 0 |
| **Keto. line** | **57***** | 0 | **57***** |

B

| MG | Plastid area ($\mu m^2$) | Number of PG | total PG area | Plastid/PG area | Grey value (colour intensity): Average PG | Grey value (colour intensity): Background | Grey value (colour intensity): Ratio |
|---|---|---|---|---|---|---|---|
| **Control** | 5.5 ± 1.9 | 16 ± 7 | 0.2 ± 0.2 | 49.4 ± 39.3 | 78.1 ± 19.6 | 189.9 ± 15.2 | 2.5 ± 0.5 |
| **β-caro. line** | **4.1 ± 1.5*** | 19 ± 12 | 0.1 ± 0.1 | 42.8 ± 19.8 | **111.1 ± 27.5**** | 189.8 ± 23.7 | **1.8 ± 0.3***** |
| **Keto. line** | 4.9 ± 2.2 | 17 ± 10 | 0.2 ± 0.2 | 30.9 ± 14.9 | **61.7 ± 13.9*** | 175.8 ± 18.2 | 3.0 ± 0.7 |

| RIPE | Plastid area ($\mu m^2$) | Number of PG | total PG area | Plastid/PG area | Grey value (colour intensity): Average PG | Grey value (colour intensity): Background | Grey value (colour intensity): Ratio |
|---|---|---|---|---|---|---|---|

| | | | | | | | | | | | | | | | | | | | | | |
|---|---|---|---|---|---|---|---|---|---|---|---|---|---|---|---|---|---|---|---|---|---|
| **Control** | 2.0 | ± | 0.8 | 8 | ± | 4 | 0.3 | ± | 0.1 | 8.4 | ± | 4.9 | 54.3 | ± | 11.4 | 114.0 | ± | 6.1 | 2.2 | ± | 0.4 |
| **β-caro. line** | **6.7** | ± | **2.3*** | 14 | ± | 8 | 1.2 | ± | 0.6 | 6.1 | ± | 1.1 | **108.5** | ± | **12.4***** | **144.3** | ± | **5.6*** | 1.3 | ± | 0.2 |
| **Keto. line** | **6.3** | ± | **2.6**** | 14 | ± | 6 | **1.6** | ± | **0.6**** | 4.1 | ± | 0.8 | **43.9** | ± | **6.6*** | **185.2** | ± | **17.0***** | **4.4** | ± | **0.6***** |

*Grey value range: black = 0, white = 255*

**Table S2: Electron microscopy related measurements of mature green and ripe tomato fruit electron micrographs**

(A) Visual quantification of membranous sacs in plastids (chloroplast and chromoplast). Data are presented as a percentage of presence of membranous sacs per total number of electron micrographs for each genotype. (B) Plastid (chloroplast and chromoplast) area and total plastoglobule (PG) area are presented as $\mu m^2$. Statistical differences compared to the control line (ANOVA analysis) are indicated in bold. $P < 0.05$, $P < 0.01$, and $P < 0.001$ are designated by *, **, and ***, respectively. P-values are listed in Table S13. The grey ratio represents the plastoglobule colour intensity normalised to the background (stroma) colour in the electron micrographs of the different genotypes. Grey values of average PG and background indicate the mean of grey values recorded within plastoglobules and the plastid's stroma, respectively. MG, mature green stage.

**% of µg/fraction**

| Control | F1 | F2 | F3 | F4 | F5 | F6 | F7 | F8 | F9 | F10 | F11 | F12 | F13 | F14 | F15 | F16 | F17 | F18 | F19 | F20 | F21 | F22 | F23 | F24 | F25 | F26 | F27 | F28 |
|---|---|---|---|---|---|---|---|---|---|---|---|---|---|---|---|---|---|---|---|---|---|---|---|---|---|---|---|---|
| **Lycopene** | 2.6 | 2.6 | 2.4 | 4.0 | 3.4 | 4.4 | 3.5 | 4.0 | 2.7 | 1.7 | 2.4 | 1.6 | 2.3 | 3.0 | 5.0 | 5.9 | 10.7 | 16.2 | 6.1 | 4.5 | 3.7 | 1.7 | 1.6 | 0.9 | 0.9 | 0.9 | 0.6 | 0.8 |
| **Lutein** | 3.5 | 3.5 | 3.5 | 3.5 | 3.5 | 3.5 | 3.5 | 3.5 | 3.5 | 3.5 | 3.5 | 3.5 | 3.5 | 3.5 | 3.6 | 3.8 | 4.1 | 4.1 | 3.6 | 3.5 | 3.6 | 3.5 | 3.5 | 3.5 | 3.5 | 3.5 | 3.5 | 3.5 |
| **β-Carotene** | 3.4 | 3.2 | 3.1 | 3.2 | 3.1 | 3.2 | 3.1 | 3.2 | 3.0 | 2.9 | 3.0 | 2.9 | 3.0 | 3.2 | 3.5 | 3.7 | 5.2 | 8.9 | 4.7 | 4.4 | 3.9 | 3.5 | 3.4 | 3.2 | 3.2 | 3.1 | 3.0 | 3.0 |
| **Phytofluene** | 4.9 | 3.7 | 3.0 | 3.1 | 2.7 | 2.8 | 2.7 | 2.9 | 2.7 | 2.5 | 2.8 | 2.7 | 3.0 | 3.9 | 4.9 | 5.1 | 6.5 | 7.5 | 4.9 | 4.4 | 3.9 | 3.5 | 3.0 | 2.7 | 2.6 | 2.5 | 2.3 | 2.4 |
| **Carotenoids** | 3.1 | 2.9 | 2.7 | 3.8 | 3.3 | 4.0 | 3.4 | 3.8 | 2.8 | 2.1 | 2.6 | 2.1 | 2.6 | 3.2 | 4.7 | 5.3 | 8.9 | 13.0 | 5.5 | 4.4 | 3.7 | 2.3 | 2.2 | 1.6 | 1.6 | 1.6 | 1.4 | 1.5 |
| **Cyclic carotenoids** | 3.5 | 3.4 | 3.4 | 3.4 | 3.4 | 3.4 | 3.4 | 3.4 | 3.3 | 3.3 | 3.3 | 3.3 | 3.3 | 3.4 | 3.5 | 3.8 | 4.5 | 5.8 | 4.0 | 3.8 | 3.7 | 3.5 | 3.5 | 3.4 | 3.4 | 3.4 | 3.3 | 3.3 |

| Keto line | F1 | F2 | F3 | F4 | F5 | F6 | F7 | F8 | F9 | F10 | F11 | F12 | F13 | F14 | F15 | F16 | F17 | F18 | F19 | F20 | F21 | F22 | F23 | F24 | F25 | F26 | F27 | F28 |
|---|---|---|---|---|---|---|---|---|---|---|---|---|---|---|---|---|---|---|---|---|---|---|---|---|---|---|---|---|
| **Astaxanthin** | 1.8 | 0.6 | 1.0 | 1.2 | 1.0 | 1.0 | 0.8 | 0.8 | 1.1 | 0.8 | 2.0 | 2.5 | 3.3 | 5.1 | 7.8 | 7.5 | 17.7 | 34.5 | 5.1 | 2.3 | 1.3 | 0.4 | 0.1 | 0.0 | 0.0 | 0.0 | 0.0 | 0.1 |
| **Phoenicoxanthin** | 4.3 | 1.9 | 2.0 | 1.9 | 1.5 | 1.2 | 1.2 | 1.1 | 1.6 | 0.8 | 1.5 | 1.8 | 2.4 | 3.8 | 5.7 | 6.1 | 16.9 | 34.6 | 5.0 | 2.3 | 1.3 | 0.4 | 0.2 | 0.0 | 0.0 | 0.0 | 0.0 | 0.2 |
| **Canthaxanthin** | 5.1 | 2.7 | 2.7 | 2.5 | 2.2 | 1.6 | 1.7 | 1.6 | 2.1 | 1.2 | 1.6 | 1.8 | 2.3 | 3.4 | 5.0 | 5.5 | 14.8 | 27.7 | 5.2 | 2.9 | 1.9 | 1.1 | 0.8 | 0.5 | 0.5 | 0.4 | 0.5 | 0.9 |
| **3'OH-Echinenone** | 5.3 | 3.0 | 3.2 | 3.2 | 2.6 | 2.1 | 2.4 | 2.2 | 2.3 | 1.9 | 2.1 | 2.1 | 2.6 | 3.6 | 5.1 | 4.7 | 11.9 | 23.9 | 4.0 | 2.4 | 1.9 | 1.3 | 1.2 | 1.0 | 1.0 | 0.9 | 0.9 | 1.2 |
| **3OH-Echinenone** | 2.7 | 2.6 | 3.9 | 3.7 | 3.4 | 2.9 | 2.9 | 2.7 | 3.4 | 2.8 | 2.5 | 2.8 | 3.1 | 3.6 | 4.1 | 4.3 | 8.5 | 15.4 | 4.8 | 3.7 | 2.3 | 2.3 | 2.1 | 2.0 | 2.0 | 2.0 | 1.9 | 2.0 |
| **Echinenone** | 3.6 | 2.6 | 2.6 | 2.7 | 2.7 | 2.1 | 2.3 | 2.2 | 2.6 | 1.9 | 2.3 | 2.5 | 2.8 | 3.4 | 4.5 | 4.3 | 12.5 | 24.4 | 4.4 | 2.6 | 1.9 | 1.5 | 1.4 | 1.2 | 1.2 | 1.2 | 1.2 | 1.4 |
| **Phoenicoxanthin-C14:0** | 5.1 | 2.2 | 2.1 | 1.9 | 1.4 | 0.7 | 0.8 | 0.8 | 1.3 | 0.3 | 0.7 | 1.0 | 1.8 | 3.1 | 5.3 | 5.5 | 17.0 | 39.5 | 5.4 | 2.4 | 1.2 | 0.2 | 0.1 | 0.0 | 0.0 | 0.0 | 0.0 | 0.0 |
| **Adonixanthin-C14:1** | 5.0 | 2.7 | 2.9 | 2.6 | 2.3 | 1.6 | 1.9 | 1.8 | 2.2 | 1.6 | 1.7 | 1.9 | 2.5 | 3.5 | 4.9 | 4.7 | 13.0 | 28.6 | 4.7 | 2.3 | 1.8 | 1.0 | 0.9 | 0.7 | 0.7 | 0.6 | 1.3 | 0.9 |
| **Phoenicoxanthin-C16:0** | 5.5 | 2.5 | 2.4 | 2.1 | 1.7 | 0.8 | 0.9 | 0.9 | 1.5 | 0.1 | 0.6 | 1.0 | 1.6 | 2.9 | 5.0 | 5.3 | 17.7 | 38.4 | 5.8 | 2.3 | 1.0 | 0.1 | 0.0 | 0.0 | 0.0 | 0.0 | 0.0 | 0.0 |
| **Adonixanthin-C16:1** | 5.0 | 2.6 | 2.8 | 2.8 | 2.4 | 1.5 | 1.8 | 1.9 | 2.1 | 1.4 | 1.8 | 1.8 | 2.3 | 3.1 | 4.5 | 4.1 | 10.9 | 24.5 | 9.7 | 6.5 | 1.6 | 1.1 | 0.8 | 0.5 | 0.6 | 0.5 | 0.5 | 0.8 |
| **Adonixanthin epoxide** | 3.5 | 3.4 | 3.4 | 3.4 | 3.4 | 3.4 | 3.4 | 3.4 | 3.4 | 3.4 | 3.5 | 3.5 | 3.5 | 3.7 | 3.9 | 3.9 | 4.6 | 5.5 | 3.7 | 3.5 | 3.4 | 3.4 | 3.3 | 3.3 | 3.3 | 3.3 | 3.3 | 3.3 |
| **Free ketocarotenoids** | 4.7 | 2.5 | 2.6 | 2.5 | 2.1 | 1.6 | 1.7 | 1.6 | 2.0 | 1.3 | 1.7 | 1.9 | 2.4 | 3.5 | 5.2 | 5.5 | 14.8 | 28.4 | 5.0 | 2.7 | 1.8 | 1.0 | 0.8 | 0.5 | 0.5 | 0.4 | 0.5 | 0.8 |

| | | | | | | | | | | | | | | | | | | | | | | | | | | | | |
|---|---|---|---|---|---|---|---|---|---|---|---|---|---|---|---|---|---|---|---|---|---|---|---|---|---|---|---|---|
| **Ketocartenoid esters** | 5.2 | 2.4 | 2.4 | 2.2 | 1.8 | 0.9 | 1.1 | 1.1 | 1.6 | 0.5 | 0.9 | 1.2 | 1.9 | 3.1 | 5.0 | 5.2 | 15.9 | 35.7 | 6.0 | 2.9 | 1.3 | 0.4 | 0.3 | 0.2 | 0.2 | 0.2 | 0.2 | 0.2 |
| **β-Carotene** | 2.2 | 2.4 | 2.5 | 2.7 | 2.8 | 2.2 | 2.3 | 2.2 | 2.4 | 2.0 | 2.3 | 2.5 | 2.9 | 3.5 | 4.5 | 4.0 | 12.4 | 24.0 | 4.4 | 3.1 | 1.9 | 1.8 | 1.6 | 1.5 | 1.5 | 1.4 | 1.7 | 1.5 |

| β-caro line | F1 | F2 | F3 | F4 | F5 | F6 | F7 | F8 | F9 | F10 | F11 | F12 | F13 | F14 | F15 | F16 | F17 | F18 | F19 | F20 | F21 | F22 | F23 | F24 | F25 | F26 | F27 | F28 |
|---|---|---|---|---|---|---|---|---|---|---|---|---|---|---|---|---|---|---|---|---|---|---|---|---|---|---|---|---|
| **Neo/Violaxanthin** | 0.0 | 0.0 | 0.0 | 0.0 | 3.8 | 3.8 | 3.8 | 3.9 | 3.9 | 3.9 | 3.9 | 4.0 | 4.2 | 4.3 | 4.9 | 5.0 | 4.9 | 5.6 | 4.5 | 4.2 | 3.9 | 4.0 | 3.9 | 3.9 | 3.9 | 3.9 | 3.9 | 4.0 |
| **Lutein** | 3.5 | 3.5 | 3.5 | 3.5 | 3.5 | 3.5 | 3.5 | 3.5 | 3.5 | 3.5 | 3.5 | 3.5 | 3.6 | 3.6 | 3.8 | 3.8 | 3.9 | 4.2 | 3.8 | 3.6 | 3.5 | 3.5 | 3.5 | 3.5 | 3.5 | 3.5 | 3.5 | 3.5 |
| **Lycopene** | 0.0 | 0.0 | 0.0 | 3.4 | 3.5 | 3.8 | 3.9 | 3.8 | 3.7 | 3.6 | 3.7 | 3.7 | 3.7 | 3.7 | 4.0 | 4.1 | 4.8 | 10.6 | 5.4 | 3.7 | 3.5 | 3.4 | 3.4 | 3.3 | 3.3 | 3.3 | 3.3 | 3.3 |
| **α-Carotene** | 3.4 | 3.4 | 3.4 | 3.5 | 3.5 | 3.6 | 3.6 | 3.7 | 3.6 | 3.6 | 3.6 | 3.6 | 3.6 | 3.6 | 3.6 | 3.6 | 3.8 | 5.0 | 3.9 | 3.5 | 3.4 | 3.4 | 3.4 | 3.4 | 3.3 | 3.3 | 3.3 | 3.4 |
| **γ-Carotene** | 0.0 | 3.3 | 3.6 | 3.6 | 3.5 | 3.5 | 3.4 | 3.3 | 3.3 | 3.3 | 3.4 | 3.2 | 3.2 | 3.3 | 3.4 | 3.8 | 4.4 | 10.0 | 5.1 | 3.6 | 3.4 | 3.3 | 3.2 | 3.2 | 3.2 | 3.2 | 3.2 | 3.2 |
| **β-Carotene** | 1.9 | 2.4 | 4.1 | 4.3 | 3.6 | 3.5 | 3.4 | 3.1 | 2.5 | 1.9 | 2.3 | 2.3 | 2.3 | 2.2 | 3.5 | 3.7 | 5.8 | 24.3 | 8.6 | 2.9 | 2.0 | 1.8 | 1.3 | 1.3 | 1.2 | 1.1 | 1.2 | 1.6 |
| **Carotenoids** | 1.9 | 2.4 | 3.2 | 3.6 | 3.5 | 3.6 | 3.5 | 3.4 | 3.1 | 2.8 | 3.0 | 3.0 | 3.0 | 3.0 | 3.7 | 3.9 | 4.9 | 14.7 | 6.3 | 3.3 | 2.8 | 2.7 | 2.5 | 2.4 | 2.4 | 2.4 | 2.4 | 2.6 |
| **Cyclic carotenoids** | 1.9 | 2.5 | 3.5 | 3.6 | 3.6 | 3.5 | 3.4 | 3.3 | 2.9 | 2.6 | 2.9 | 2.9 | 2.9 | 2.9 | 3.7 | 3.9 | 5.1 | 16.2 | 6.7 | 3.3 | 2.7 | 2.6 | 2.3 | 2.3 | 2.2 | 2.2 | 2.2 | 2.5 |

**Table S3: Carotenoid quantification of the fruit chromoplast fractions**

Carotenoid content is presented as mean in percentage of μg/fraction. F, fraction (1 ml). These data correspond to the chromoplast fractionation experiment displayed in Figure 2b and Figure S5.

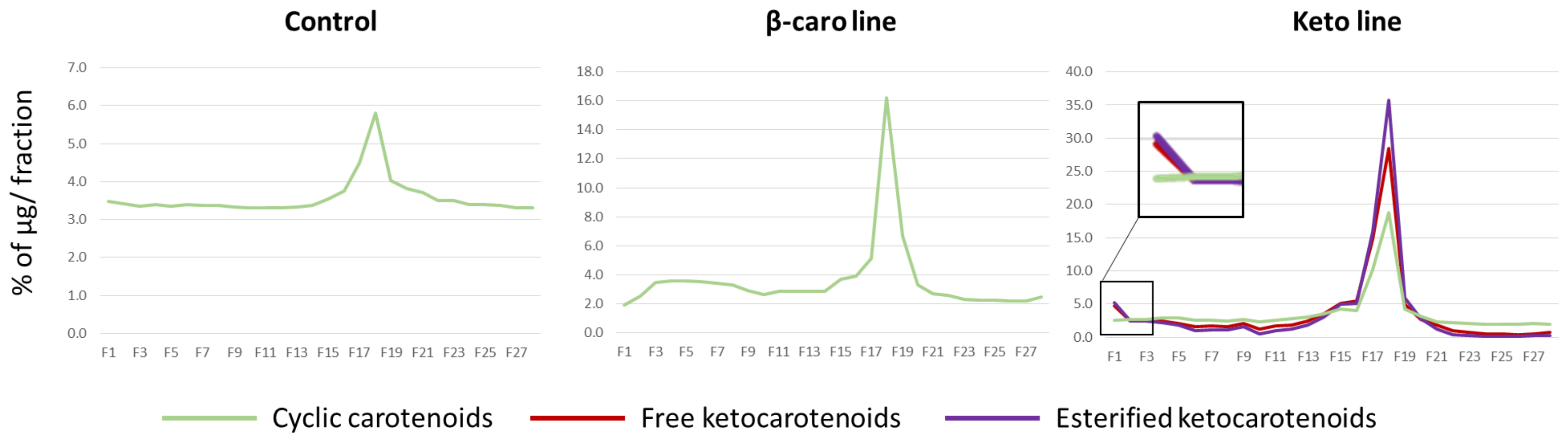


**Figure S5: Carotenoid quantification of the fruit chromoplast fractions**

Data are presented as the mean percentage of carotenoid content in a single fraction (F, 1 ml) compared with its total amount in the gradient tube (28 ml). These data correspond to the chromoplast fractionation experiment displayed in Figure 2b and Table S3.

See Excel file
**Table S4: Metabolite content in the ripe fruit**

(A) Content of all metabolites analysed in ripe fruit are presented for each biological replicate of the control, β-carotene and ketocarotenoid lines, as well as their mean in µg/g DW or arbitrary unit depending on the analytical platform. (B) List of the statistically different metabolites from the keto line vs control ripe comparison (with corresponding p-values and log2(ratio)) used to create Figure 4. (C) List of the statistically different metabolites from the β-caro line vs control ripe comparison (with corresponding p-values and log2(ratio)) used to create Figure S7.

See Excel file
**Table S5: Metabolite content in the mature green (MG) fruit**

(A) Content of all metabolites analysed in MG fruit are presented for each biological replicate of the control, β-carotene and ketocarotenoid lines, as well as their mean in µg/g DW or arbitrary unit depending on the analytical platform. (B) List of the statistically different metabolites from the keto line vs control MG comparison (with corresponding p-values and log2(ratio)) used to create Figure 4. (C) List of the statistically different metabolites from the β-caro line vs control MG comparison (with corresponding p-values and log2(ratio)) used to create Figure S7.

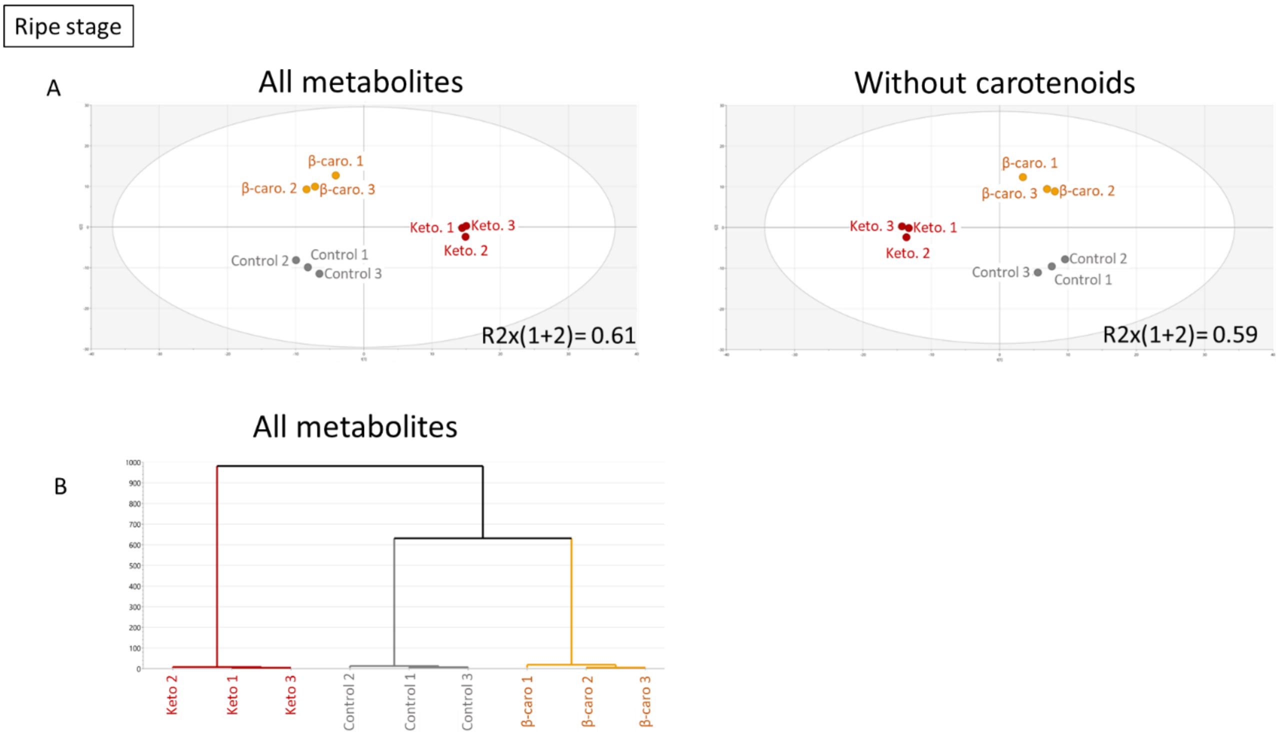


**Figure S6: Principal component analysis of all metabolites quantified in ripe fruit including or excluding the carotenoid data and metabolic hierarchical clustering of the tomato lines**

(A) Score plots of the principal component analysis of all metabolites quantified in the control, β-carotene and ketocarotenoid ripe fruit including and excluding carotenoid data (B) Hierarchical clustering of the tomato lines based on the ripe fruit metabolite data including carotenoid data.

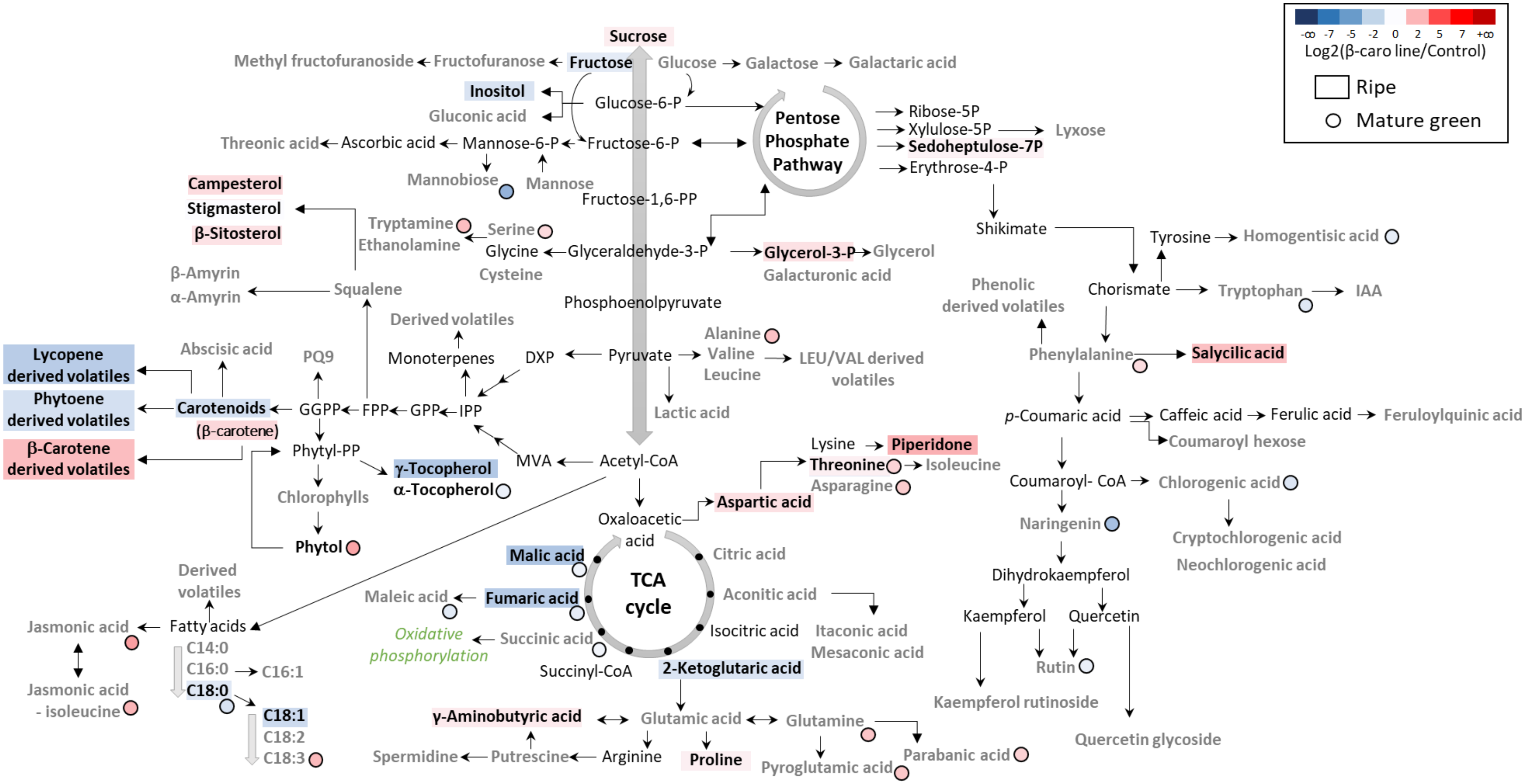


**Figure S7: Biochemical pathways displaying the fold change metabolite levels in ripe and mature green fruit of the β-carotene line compared to the control**

Data represent the log2 of the ratio of means (β-caro line/Control) and it is displayed by a colour illustrating a decrease (blue) or increase (red) of metabolite level in the β-caro line compared to the control. Compounds detected in the analytical platforms are shown in bold. The absence of significant difference is depicted by a grey bold font (t-test, $P>0.05$). Compounds showing a significant difference (black bold font) were attributed a colour representative of fold change. Metabolite levels quantified in ripe and mature green fruits are represented by a rectangle and a circle, respectively. The statistical significance of the volatile data in this display corresponds to the total content per class of compounds. The individual differences within the class of compounds are not shown. Metabolite data were compiled in Table S4 and S5. PQ9, plastoquinone 9; IAA, Indole-3-acetic acid.

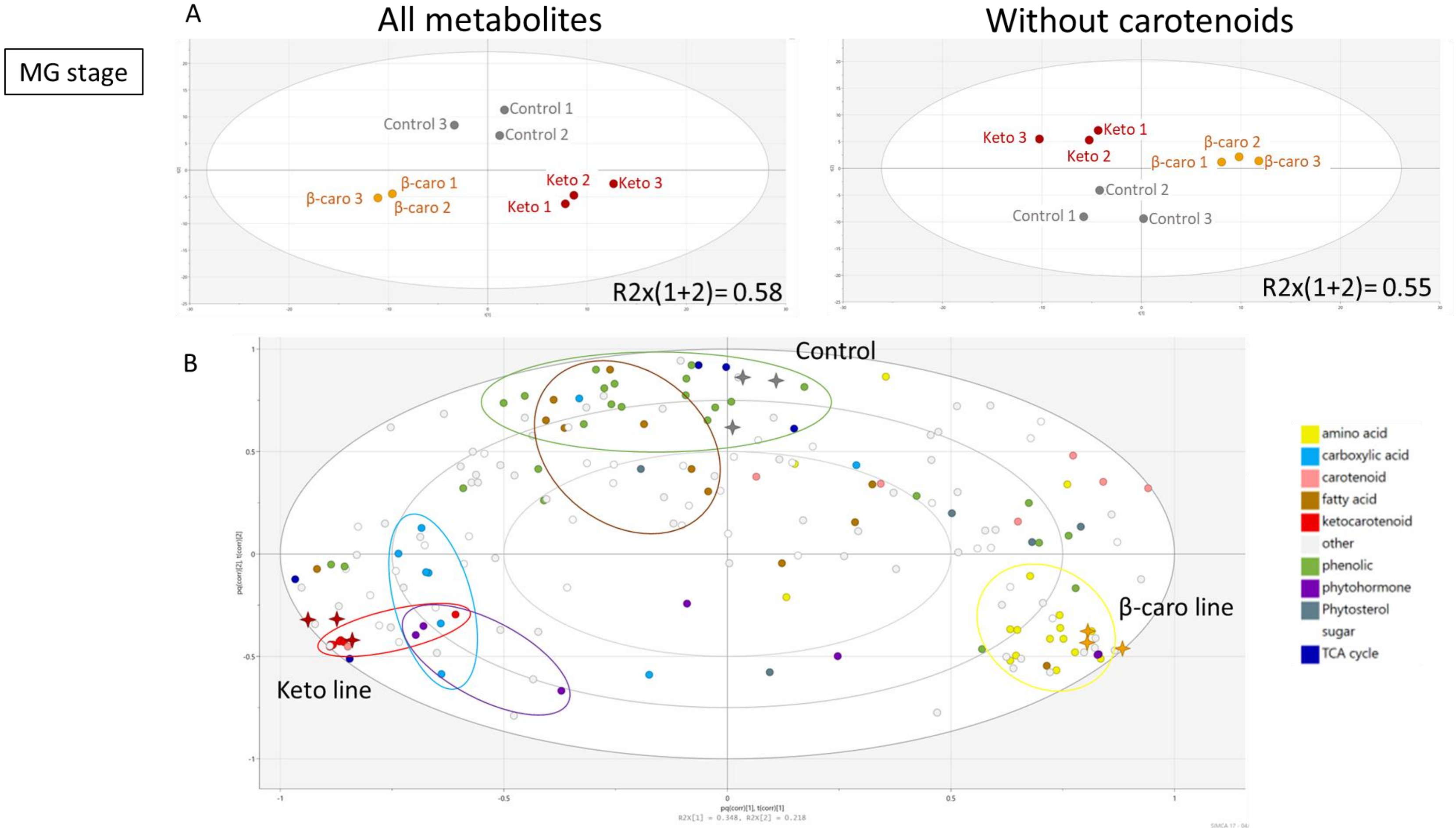


**Figure S8: Principal component analysis of all metabolites quantified in mature green fruit including or excluding the carotenoid data**

(A) Score plots of the principal component analysis of all metabolites quantified in the control, β-carotene line and ketocarotenoid line mature green fruit including and excluding the carotenoid data. (B) Loading plot of the principal component analysis of all metabolites quantified in the control, β-carotene and ketocarotenoid mature green fruit including the carotenoid data.

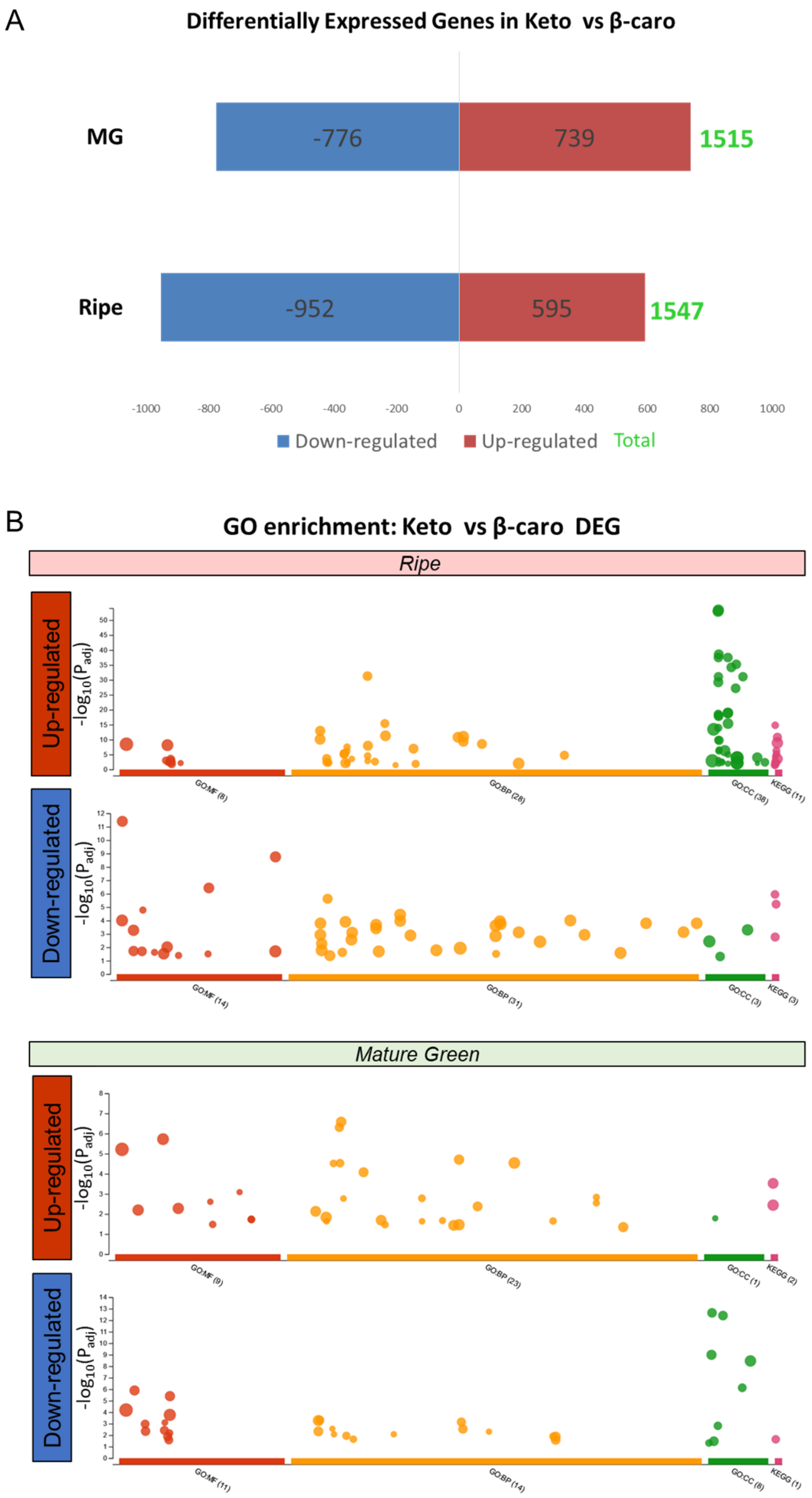


**Figure S9: Differentially expressed genes and enrichment analysis of the β-carotene/Control comparison**

(A) Up-regulated and down-regulated differentially expressed genes (DEG) count of the β-carotene/control comparison at mature green (MG) and ripe fruit stage. (B) Enriched GO terms of the β-carotene/control up-regulated and down-regulated DEGs. MF, molecular function; BP, biological process; CC, cellular component; KEGG, KEGG pathway. The full list of enriched GO terms can be found in Table S6. Gene enrichment analysis was carried out with g:Profiler (https://biit.cs.ut.ee/gprofiler/).

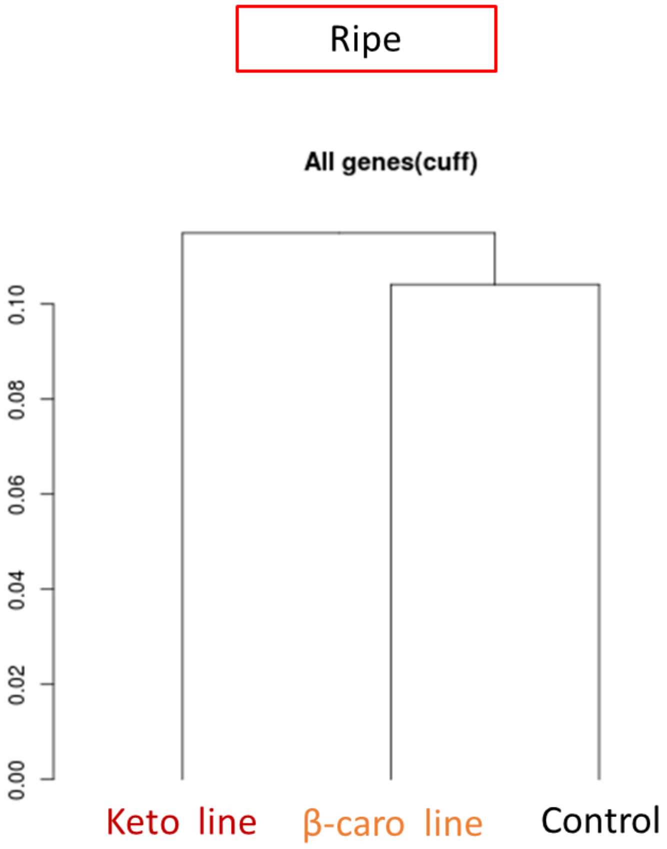


**Figure S10: Hierarchical clustering of the control, β-carotene and ketocarotenoid line RNAseq data**

Hierarchical clustering of the tomato lines based on the ripe fruit DEGs from the RNAseq data.

See Excel file

**Table S6: Gene Ontology enrichment analysis lists**

Details of Gene Ontology (GO) enrichment analyses of the RNAseq data are described in this table (DEG set studied, GO category, term name, term id, adjusted p-value, negative log10 of adjusted p-value, term size, query size, intersection size, effective domain size, intersections). The following datasets were used: Keto/Control DEGS, β-carotene/control DEGS, Keto/ β-carotene DEGS separated as up and down-regulated DEG sets and for MG and Ripe fruit stages as well as Ripe/MG DEGS for the control, β-caro line and keto line, respectively. MF, molecular function; BP, biological process; CC, cellular component; KEGG, KEGG pathway. Gene enrichment analysis was carried out with g:Profiler (https://biit.cs.ut.ee/gprofiler/).

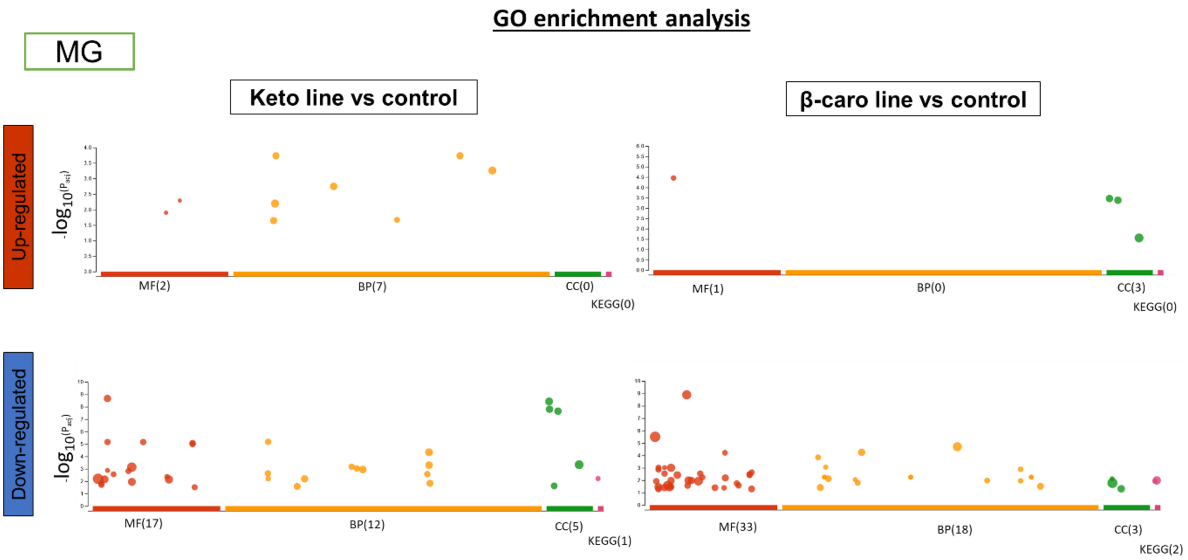


**Figure S11: Gene ontology enrichment analysis of the Keto/Control and β-carotene/Control comparison at MG**

Enriched GO terms of the Keto/control and β-carotene/control up-regulated and down-regulated DEGs in MG fruit. MF, molecular function; BP, biological process; CC, cellular component; KEGG, KEGG pathway. Gene enrichment analysis was carried out with g:Profiler (https://biit.cs.ut.ee/gprofiler/). The full list of enriched GO terms can be found in Table S6.

See Excel file

**Table S7: List of RNAseq DEGs used to create pathway visualization displays**

Details of the genes and their log2(fold change), used to create Figure 6 and Figure S12, are listed in this table.

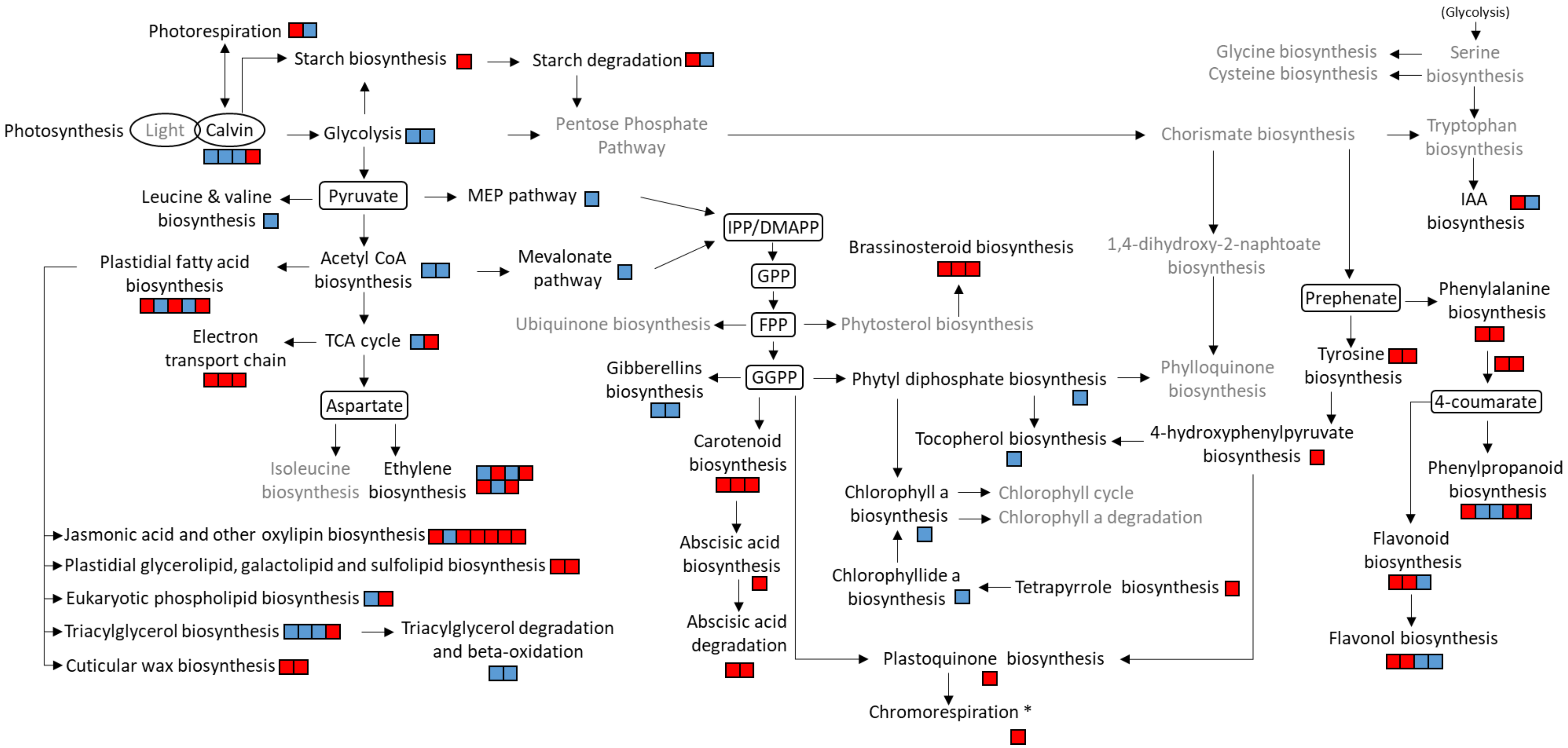


**Figure S12: Transcriptional changes in the metabolic pathways of the ripe fruit of the β-carotene line compared to the control.**

RNAseq data was used to create this display. Each square represents a gene of the referred pathway. Red and blue colours illustrate an increase and decrease in expression, respectively (positive or negative log2(Fold change)). The grey font denotes a lack of significant difference in expression level for all the genes investigated in the respective pathway (P > 0.05). All genes used to construct this figure are listed in Table S7 together with their log2(Fold change). Gramene database (www.gramene.org) as well as literature were used to compose the different metabolic pathways. IPP, Isopentenyl pyrophosphate; DMAPP, Dimethylallyl pyrophosphate; GPP, Geranyl pyrophosphate; FPP, Farnesyl diphosphate;

GGPP, Geranylgeranyl pyrophosphate ; IAA, Indole-3-acetic acid ; TCA, Tricarboxylic acid cycle ; * according to models proposed by Renato et al., 2014 and Grabsztunowicz et al., 2019.

**Plastidial redox regulatory genes_ Keto/Control**

| | | Log2(fold change) | Fold change |
|---|---|---|---|
| *Solyc04g057980* | NAD(P)H-quinone oxidoreductase subunit M, chloroplastic | 4.3 | 19.7 |
| *Solyc09g083190* | Photosynthetic NDH subunit of subcomplex B 5, chloroplastic | 2.9 | 7.5 |
| *Solyc05g026550* | NAD(P)H-quinone oxidoreductase subunit L, chloroplastic | 2.3 | 4.9 |
| *Solyc05g007780* | Photosynthetic NDH subcomplex L 2 | 1.8 | 3.5 |
| *Solyc08g081690* | NADPH oxidase | 1.4 | 2.6 |
| *Solyc09g083150* | NAD(P)H-quinone oxidoreductase subunit N, chloroplastic | 1.2 | 2.3 |
| *Solyc01g096240* | NADH-plastoquinone oxidoreductase subunit 5 (chloroplast) | 0.7 | 1.6 |
| *Solyc09g091070* | Malate dehydrogenase, chloroplastic | 0.7 | 1.6 |
| *Solyc10g082030* | 2-Cys peroxiredoxin 1 | 0.7 | 1.6 |
| *Solyc12g005630* | Cytochrome b6-f complex iron-sulfur subunit, chloroplastic | 1.1 | 2.1 |
| *Solyc08g080050* | PGR5-like protein 1A, chloroplastic | 0.8 | 1.7 |
| *Solyc02g079750* | Probable NAD(P)H dehydrogenase (quinone) FQR1-like 2 | 0.7 | 1.6 |
| *Solyc03g034140* | Probable NAD(P)H dehydrogenase (quinone) FQR1-like 3 | -1.9 | 0.3 |
| *Solyc03g115005* | PsbQ-like protein 3, chloroplastic | 3.9 | 14.9 |
| *Solyc12g005630* | Cytochrome b6-f complex iron-sulfur subunit, chloroplastic | 1.1 | 2.1 |
| *Solyc12g009400* | Pyruvate dehydrogenase E1 component subunit alpha-3, chloroplastic | 0.8 | 1.7 |
| *Solyc02g083810* | Ferredoxin--NADP reductase, leaf-type isozyme, chloroplastic | 1.2 | 2.3 |
| *Solyc09g007190* | Thioredoxin-like protein AAED1, chloroplastic isoform X1 | 1.0 | 2 |
| *Solyc03g115870* | Thioredoxin-like 1-2, chloroplastic | -1.0 | 0.5 |
| *Solyc04g078910* | Thioredoxin-like 4, chloroplastic | 1.5 | 2.8 |

**Table S8: Expression data of plastidial redox regulatory genes**

The list of genes and their expression data (fold change) were retrieved from the RNA seq dataset.

See Excel file

**Table S9: Gene-metabolite correlation matrix of the ripe keto line / control comparison**

Pearson correlation analysis between the statistically different metabolites found in the keto/control comparison (p-value $< 0.05$) and DEGs of the keto/control comparison involved in biosynthetic pathways (List of genes used in Figure 6 plus genes that mapped in Mapman (https://mapman.gabipd.org/)) was performed. Metabolite and gene expression levels were expressed as log2(fold change = keto/control) for this analysis. The correlation values obtained as well as the list of metabolites and genes included in the analysis can be found in this table.

See Excel file

**Table S10: Gene enrichment analysis of the down-regulated Ripe/MG DEGs of the keto line versus the control line**

Fisher's Exact Test was carried out to obtain the enriched GO terms with the OmicsBox 1.4.11 software.

See Excel file

**Table S11: Gene enrichment analysis of the up-regulated DEGs of the keto/control versus β-caro/control comparison in ripe fruit**

Fisher's Exact Test was carried out to obtain the enriched GO terms with the OmicsBox 1.4.11 software. This table is related to Figure 5C.

See Excel file

**Table S12: Gene enrichment analysis of the common DEGs from the Keto/control and Keto/β-caro sets in ripe fruit**

Gene enrichment analysis was carried out with g:Profiler (https://biit.cs.ut.ee/gprofiler/). MF, molecular function; BP, biological process; CC, cellular component; KEGG, KEGG pathway.

See Excel file

**Table S13: Statistical analysis**

Details of the statistical analyses performed in SPPS software are described in this table.

See Excel file

**Table S14: Metabolite identification features**

Identification features for all metabolites analysed in this study are described in this table.
